# Neutrophil extracellular traps formation is associated with postoperative complications in neonates and infants undergoing congenital cardiac surgery

**DOI:** 10.1101/2023.12.21.572768

**Authors:** Wiriya Maisat, Lifei Hou, Sumiti Sandhu, Yi-Cheng Sin, Samuel Kim, Hanna Van Pelt, Yue Chen, Sirisha Emani, Sek Won Kong, Sitram Emani, Juan Ibla, Koichi Yuki

**Author notes:** Correspondence: Koichi Yuki, M.D., Department of Anesthesiology, Critical Care and Pain Medicine Cardiac Anesthesia Division, Boston Children’s Hospital. Author contribution: Idea and Experiment design- W.M., L.H., Y.C., S.E., S.W.K., J.C, and K.Y. Experiment execution- W.M., L.H., S.S., Y.S., S.E. Manuscript draft- W.M., K.Y. Manuscript editing- S.K., H.V.P., S.E., Grant acquisition- K.Y.

## Abstract

Pediatric patients with congenital heart diseases (CHD) often undergo surgical repair on cardiopulmonary bypass (CPB). Despite a significant medical and surgical improvement, the mortality of neonates and infants remains high. Damage-associated molecular patterns (DAMPs) are endogenous molecules released from injured/damaged tissues as danger signals. We examined 101 pediatric patients who underwent congenital cardiac surgery on CPB. The mortality rate was 4.0%, and the complication rate was 31.6%. We found that neonates/infants experienced multiple complications most, consistent with the previous knowledge. Neonates and infants in the complication group had received more transfusion intraoperatively than the non-complication arm with lower maximum amplitude (MA) on rewarming CPB thromboelastography (TEG). Despite TEG profiles were comparable at ICU admission between the two groups, the complication arm had higher postoperative chest tube output, requiring more blood transfusion. The complication group showed greater neutrophil extracellular traps (NETs) formation at the end of CPB and postoperatively. Plasma histones and high mobility group box 1 (HMGB1) levels were significantly higher in the complication arm. Both induced NETs *in vitro* and *in vivo*. As histones and HMGB1 target Toll-like receptor (TLR)2 and TLR4, their mRNA expression in neutrophils was upregulated in the complication arm. Taken together, NETs play a major role in postoperative complication in pediatric cardiac surgery and would be considered a target for intervention.

**Key points:**

- Neonates and infants showed highest postoperative complications with more upregulation of inflammatory transcriptomes of neutrophils.
- Neonates and infants with organ dysfunction had more NETs formation with higher plasma histones and HMGB1 levels.

## Introduction

CHD is the most common live birth anomaly with the incidence of 4-10/1,000^1-3^. Although their overall mortality has significantly improved^4-6^, the outcome of neonates and infants undergoing cardiac surgery remains associated with high morbidity and mortality, caused largely by organ injury/failure (in-hospital mortality of 6.9%)^7,8^. Cardiac surgery involves extensive dissection, tissue ischemia and reperfusion during cardiopulmonary bypass (CPB) and transfusion ^9^. In complex neonatal and infant surgeries, complete circulatory arrest or regional perfusion limited to the brain is often employed, increasing the vulnerability of multiple organs to ischemia-reperfusion injury ^10,11^. Damaged cells/tissues release damage-associated molecular patterns (DAMPs) into the bloodstream^12^. The release of DAMPs can be a double-edge sword to host ^13^. On one hand, DAMPs attract cells at the site of release for tissue repair and regeneration^14^. However, when the systemic inflammatory response is exaggerated by DAMPs, organ dysfunction/failure ensues^14,15^.

A subset of DAMPs has been reported in the setting of trauma, sepsis, and adult cardiac surgery ^16-19^. They are recognized by pattern recognition receptors (PRRs) such as TLR2 and TLR4 on innate immune cells and platelets ^13^. Neutrophils are the most abundant innate immune cells and first responders to inflammation for tissue repair^20^. However, their robust effector functions can damage tissues^21^. For example, they can induce NETs and cause microthrombosis ^22-25^. Elevated levels of DAMPs have been positively correlated with systemic inflammatory response syndrome (SIRS) and multiple organ injury in trauma and sepsis patients ^19,26^. Immunological/coagulation profiles, and their relationship with postoperative outcomes remain less delineated in patients with CHDs undergoing cardiac surgery. We hypothesized that congenital cardiac surgery would induce the release of circulating DAMPs, potentially triggering neutrophil activation and an increased susceptibility to thrombotic events and organ injury, particularly in neonates and infants. To test this hypothesis, we aimed to characterize the longitudinal profiles of neutrophils, coagulation responses and DAMPs in patients undergoing congenital cardiac surgery with CPB. Furthermore, we sought to establish the association between these factors and the occurrence of postoperative organ dysfunction and thrombosis. We also examined the effect of these DAMPs *in vitro* and *in vivo*. Understanding the mechanisms of postoperative organ dysfunction can lead us to the discovery of preventive and therapeutic interventions against it in this vulnerable population.

## Methods

### Study design and setting

This single-center prospective cohort study was conducted at a quaternary academic pediatric medical center. The study was approved by the Institutional Review Board at Boston Children’s Hospital, and written informed consent was obtained from a parent or legal guardian.

## Results

### Neutrophil and monocyte counts were significantly increased postoperatively

Neutrophils and monocytes serve an essential role in initiating the immune response at the sites of tissue injury^27,28^. So far, perioperative leukocyte profiles were studied primarily in adult patients undergoing non-cardiac surgery ^29,30^. Here we profiled in pediatric patients undergoing cardiac surgery on CPB. The characteristics of 101 patients are described in **Table 1**. Average age was 22.1 months with 45% male. Neonates and infants were the most with 70 patients. We observed a modest decline in neutrophil and monocyte counts during the rewarming period (T1) compared to baseline (T0), which could be attributable, at least in part, to a dilution from CPB priming volume. Upon ICU admission (T2), we observed a significant increase in both neutrophil and monocyte counts from their baseline levels (**Fig. 1A**). At ICU admission, the neutrophil counts were approximately two times higher than the baseline levels, and by postoperative day 1 (T3), approximately three times higher. We also examined platelet counts because the coagulation component is critical in cardiac surgery. We observed a significant reduction in platelet numbers at rewarming and ICU admission, with a subsequent return to baseline levels by postoperative day 1 (**Fig. 1A**). At rewarming, platelet numbers dropped to 25% of the baseline levels.

**Fig. 1.**
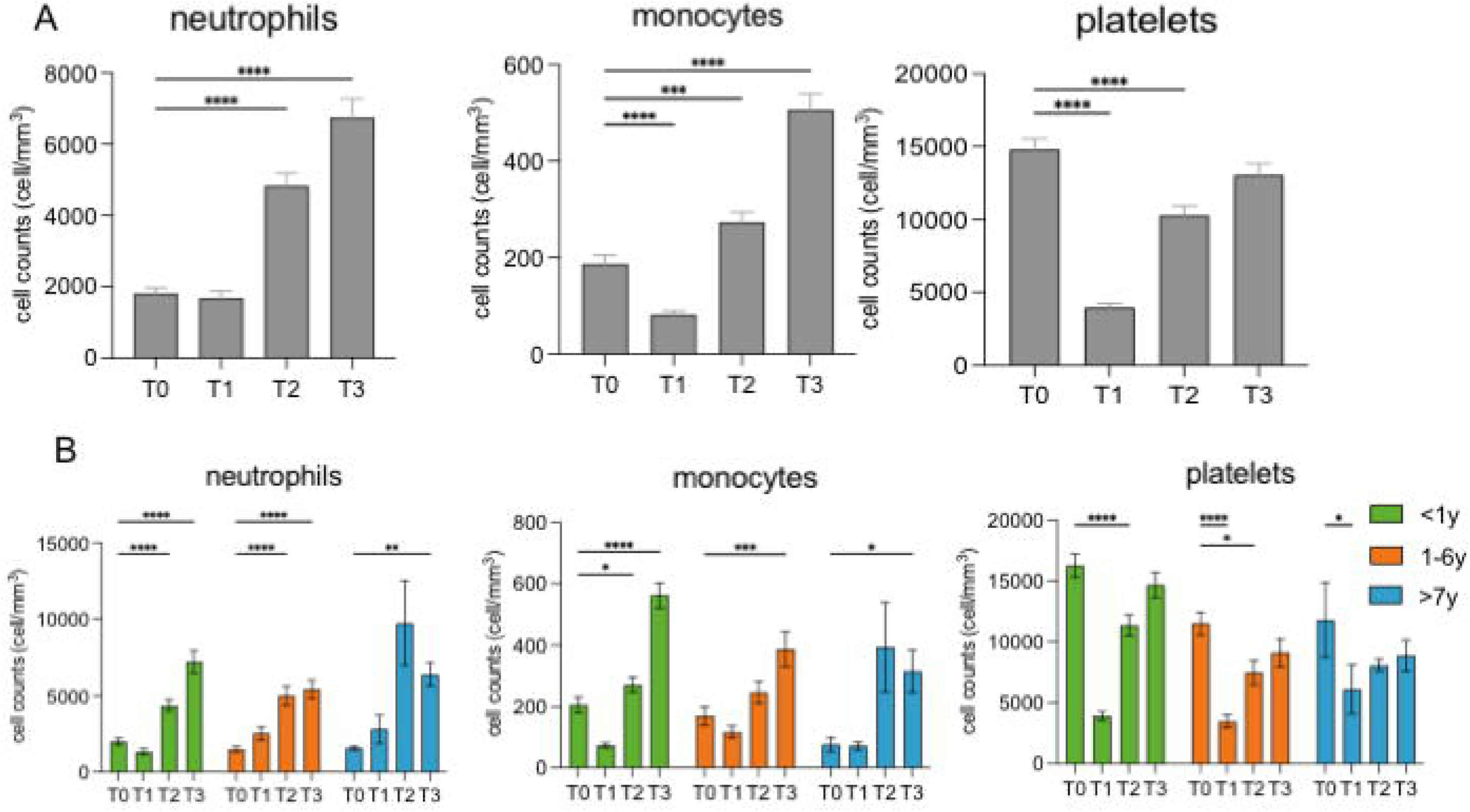
Temporal changes in neutrophil, monocyte, and platelet counts and TEG parameters following congenital cardiac surgery with CPB. **(A)** Overall neutrophil, monocyte and platelet counts from T0 to T3 in the overall cohort. **(B)** Subgroup analysis of neutrophil, monocyte, and platelet counts in patients of different age groups. (**A-B**) Data are presented as mean ± SEM. Statistical analysis was performed using two-way analysis of variance (ANOVA) with multiple comparisons test and *post hoc* Dunnett’s correction. *, **, ***, and **** denote p<0.05, 0.01, 0.001, and 0.0001, respectively. **(C)** TEG parameters including l1-angle, reaction (R) time, kinetic (k) time, and maximum amplitude (MA) values from T0 to T3 in the overall cohort (left to right). **(D)** Subgroup analysis of TEG parameters in patients of different age groups. (**C-D**) Data are presented as median and interquartile range (IQR). Statistical analysis was performed using two-way analysis of variance (ANOVA) with multiple comparisons test and *post hoc* Dunnett’s correction. *, **, ***, and **** denote p<0.05, 0.01, 0.001, and 0.0001, respectively.

**Table 1.**
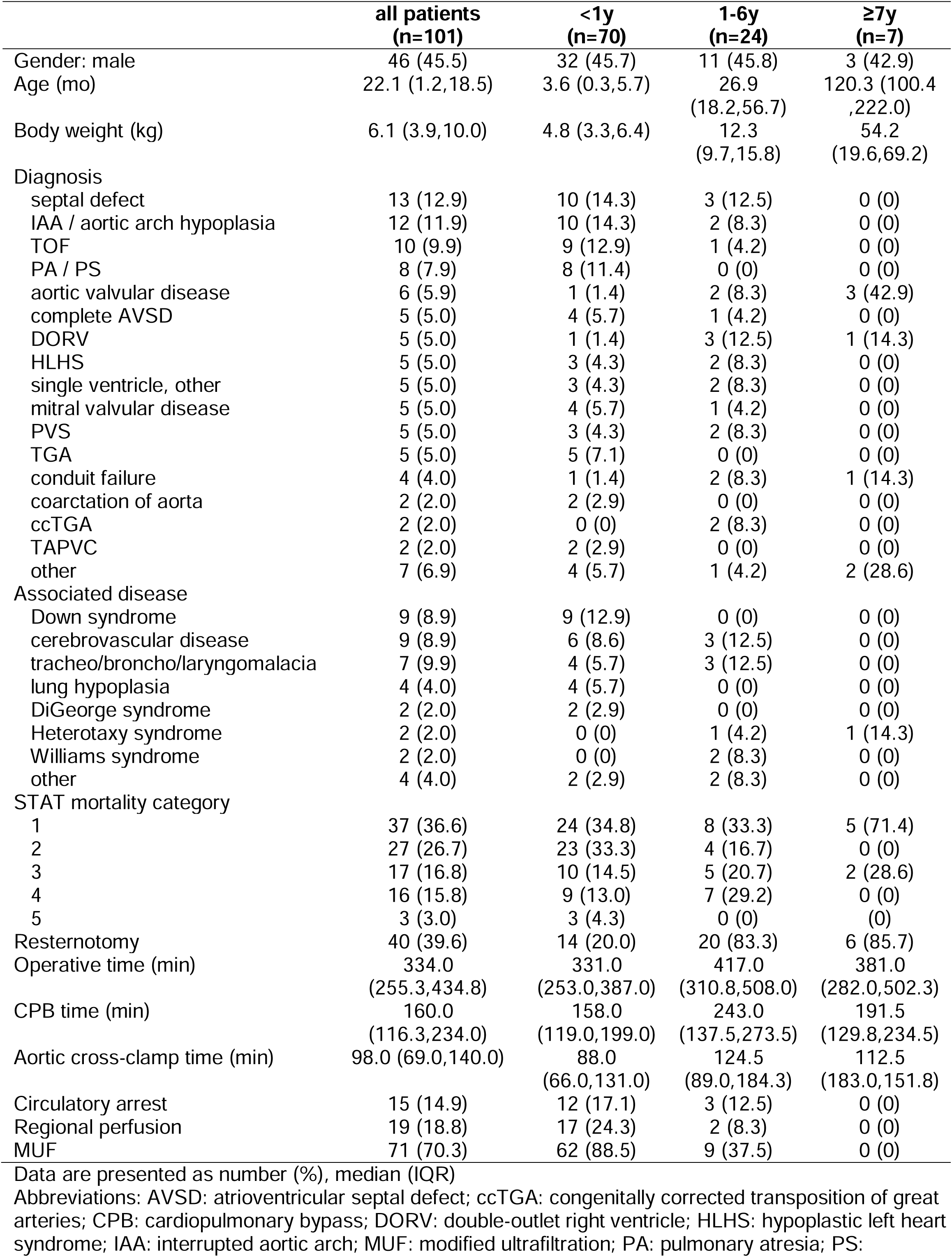
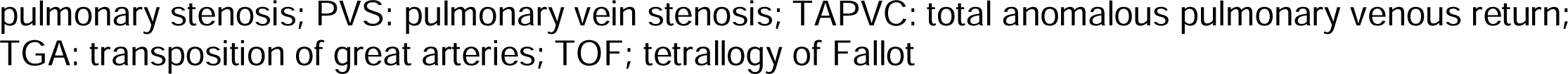
Patient characteristics.

|  | **all patients**<br>**(n=101)** | **<1y**<br>**(n=70)** | **1-6y**<br>**(n=24)** | **≥7y**<br>**(n=7)** |
| --- | --- | --- | --- | --- |
| Gender: male | 46 (45.5) | 32 (45.7) | 11 (45.8) | 3 (42.9) |
| Age (mo) | 22.1 (1.2,18.5) | 3.6 (0.3,5.7) | 26.9<br>(18.2,56.7) | 120.3 (100.4<br>,222.0) |
| Body weight (kg) | 6.1 (3.9,10.0) | 4.8 (3.3,6.4) | 12.3<br>(9.7,15.8) | 54.2<br>(19.6,69.2) |
| Diagnosis |  |  |  |  |
| septal defect | 13 (12.9) | 10 (14.3) | 3 (12.5) | 0 (0) |
| IAA / aortic arch hypoplasia | 12 (11.9) | 10 (14.3) | 2 (8.3) | 0 (0) |
| TOF | 10 (9.9) | 9 (12.9) | 1 (4.2) | 0 (0) |
| PA / PS | 8 (7.9) | 8 (11.4) | 0 (0) | 0 (0) |
| aortic valvular disease | 6 (5.9) | 1 (1.4) | 2 (8.3) | 3 (42.9) |
| complete AVSD | 5 (5.0) | 4 (5.7) | 1 (4.2) | 0 (0) |
| DORV | 5 (5.0) | 1 (1.4) | 3 (12.5) | 1 (14.3) |
| HLHS | 5 (5.0) | 3 (4.3) | 2 (8.3) | 0 (0) |
| single ventricle, other | 5 (5.0) | 3 (4.3) | 2 (8.3) | 0 (0) |
| mitral valvular disease | 5 (5.0) | 4 (5.7) | 1 (4.2) | 0 (0) |
| PVS | 5 (5.0) | 3 (4.3) | 2 (8.3) | 0 (0) |
| TGA | 5 (5.0) | 5 (7.1) | 0 (0) | 0 (0) |
| conduit failure | 4 (4.0) | 1 (1.4) | 2 (8.3) | 1 (14.3) |
| coarctation of aorta | 2 (2.0) | 2 (2.9) | 0 (0) | 0 (0) |
| ccTGA | 2 (2.0) | 0 (0) | 2 (8.3) | 0 (0) |
| TAPVC | 2 (2.0) | 2 (2.9) | 0 (0) | 0 (0) |
| other | 7 (6.9) | 4 (5.7) | 1 (4.2) | 2 (28.6) |
| Associated disease |  |  |  |  |
| Down syndrome | 9 (8.9) | 9 (12.9) | 0 (0) | 0 (0) |
| cerebrovascular disease | 9 (8.9) | 6 (8.6) | 3 (12.5) | 0 (0) |
| tracheo/broncho/laryngomalacia | 7 (9.9) | 4 (5.7) | 3 (12.5) | 0 (0) |
| lung hypoplasia | 4 (4.0) | 4 (5.7) | 0 (0) | 0 (0) |
| DiGeorge syndrome | 2 (2.0) | 2 (2.9) | 0 (0) | 0 (0) |
| Heterotaxy syndrome | 2 (2.0) | 0 (0) | 1 (4.2) | 1 (14.3) |
| Williams syndrome | 2 (2.0) | 0 (0) | 2 (8.3) | 0 (0) |
| other | 4 (4.0) | 2 (2.9) | 2 (8.3) | 0 (0) |
| STAT mortality category |  |  |  |  |
| 1 | 37 (36.6) | 24 (34.8) | 8 (33.3) | 5 (71.4) |
| 2 | 27 (26.7) | 23 (33.3) | 4 (16.7) | 0 (0) |
| 3 | 17 (16.8) | 10 (14.5) | 5 (20.7) | 2 (28.6) |
| 4 | 16 (15.8) | 9 (13.0) | 7 (29.2) | 0 (0) |
| 5 | 3 (3.0) | 3 (4.3) | 0 (0) | 0 (0) |
| Resternotomy | 40 (39.6) | 14 (20.0) | 20 (83.3) | 6 (85.7) |
| Operative time (min) | 334.0<br>(255.3,434.8) | 331.0<br>(253.0,387.0) | 417.0<br>(310.8,508.0) | 381.0<br>(282.0,502.3) |
| CPB time (min) | 160.0<br>(116.3,234.0) | 158.0<br>(119.0,199.0) | 243.0<br>(137.5,273.5) | 191.5<br>(129.8,234.5) |
| Aortic cross-clamp time (min) | 98.0 (69.0,140.0) | 88.0<br>(66.0,131.0) | 124.5<br>(89.0,184.3) | 112.5<br>(183.0,151.8) |
| Circulatory arrest | 15 (14.9) | 12 (17.1) | 3 (12.5) | 0 (0) |
| Regional perfusion | 19 (18.8) | 17 (24.3) | 2 (8.3) | 0 (0) |
| MUF | 71 (70.3) | 62 (88.5) | 9 (37.5) | 0 (0) |

Data are presented as number (%), median (IQR)
Abbreviations: AVSD: atrioventricular septal defect; ccTGA: congenitally corrected transposition of great arteries; CPB: cardiopulmonary bypass; DORV: double-outlet right ventricle; HLHS: hypoplastic left heart syndrome; IAA: interrupted aortic arch; MUF: modified ultrafiltration; PA: pulmonary atresia; PS:
pulmonary stenosis; PVS: pulmonary vein stenosis; TAPVC: total anomalous pulmonary venous return; TGA: transposition of great arteries; TOF; tetralogy of Fallot

### Neutrophil and monocyte counts in younger patients peaked on postoperative day 1

We also performed a subgroup analysis of neutrophil, monocyte and platelet counts by dividing into neonates/infants, patients aged 1-6 years, and patients older than 7 years. There was a tendency toward a reduction of neutrophil and monocyte counts at postoperative day 1 compared to ICU admission in patients older than 7 years (**Fig. 1B**). In contrast, the neonates/infants and age 1-6 groups exhibited a consistent increase in neutrophil and monocyte counts until postoperative day 1 (**Fig. 1B**). These results suggest that immunological responses to cardiac surgery on CPB may differ based on the age of the patients. In contrast, the trend of platelet counts was comparable among the three groups.

### The use of CPB caused a hypocoagulable state at the rewarming CPB phase

We also examined TEG parameters to assess the coagulation status. Our data revealed distinct TEG value patterns (**Fig. 1C**). The k time and reaction (R) time exhibited a prolonged trend, reaching their peak levels at the rewarming phase. Subsequently, these parameters gradually declined but remained significantly prolonged compared to baseline levels until postoperative day 1. In parallel, the l1-angle and maximum amplitude (MA) consistently significantly declined during the rewarming phase and continued to exhibit lower values until postoperative day 1. These findings collectively indicate a hypocoagulable state, associated with delayed clot initiation, impaired fibrin formation, and reduced clot strength (**Fig. 1C**). Upon subgroup analysis, we also observed a similar pattern in neonates/infants and age 1-6 groups, suggesting that these age groups were more affected by CPB-induced changes in general coagulation status (**Fig. 1D**). In contrast, older patients (≥ 7 years) appeared to be less affected by these alterations, different from platelet count data.

### RNA sequencing analysis revealed neutrophil activation following CPB

The increase in neutrophil counts after CPB is a physiological response to the surgical stress and associated inflammatory stimuli^31^. However, excessive or prolonged activation of neutrophils potentially contributes to complications such as organ dysfunction and tissue damage ^32^. To investigate neutrophil activation, we conducted RNA sequencing analysis of neutrophils at baseline and ICU admission. We detected a total of 56,131 genes. We identified 1,512 upregulated and 1,458 downregulated differentially expressed genes (DEGs) (**Fig. 2A**). We performed gene ontology (GO) enrichment analysis. Among the top 20 enriched GO terms associated with upregulated DEGs, we observed a significant involvement in various inflammatory processes such as innate immune response, apoptosis, and regulation of nuclear factor κB (NFKB) (**Fig. 2B**). The Kyoto Encyclopedia of Genes and Genomes (KEGG) pathway analysis revealed that the upregulated DEGs were associated with several inflammatory signaling pathways and apoptosis (**Fig. 2C**). These pathways included phosphatidylinositol 3’- kinase (PI3K)-Akt, mitogen-activated protein kinase (MAPK), Ras-associated protein-1 (Rap 1), hypoxia-inducible factor (HIF)-1 and tumor necrosis factor (TNF) signaling pathways. These findings collectively support the presence of intense pro-inflammatory responses and cellular stress that neutrophils experienced in the early postoperative period.

**Fig. 2.**
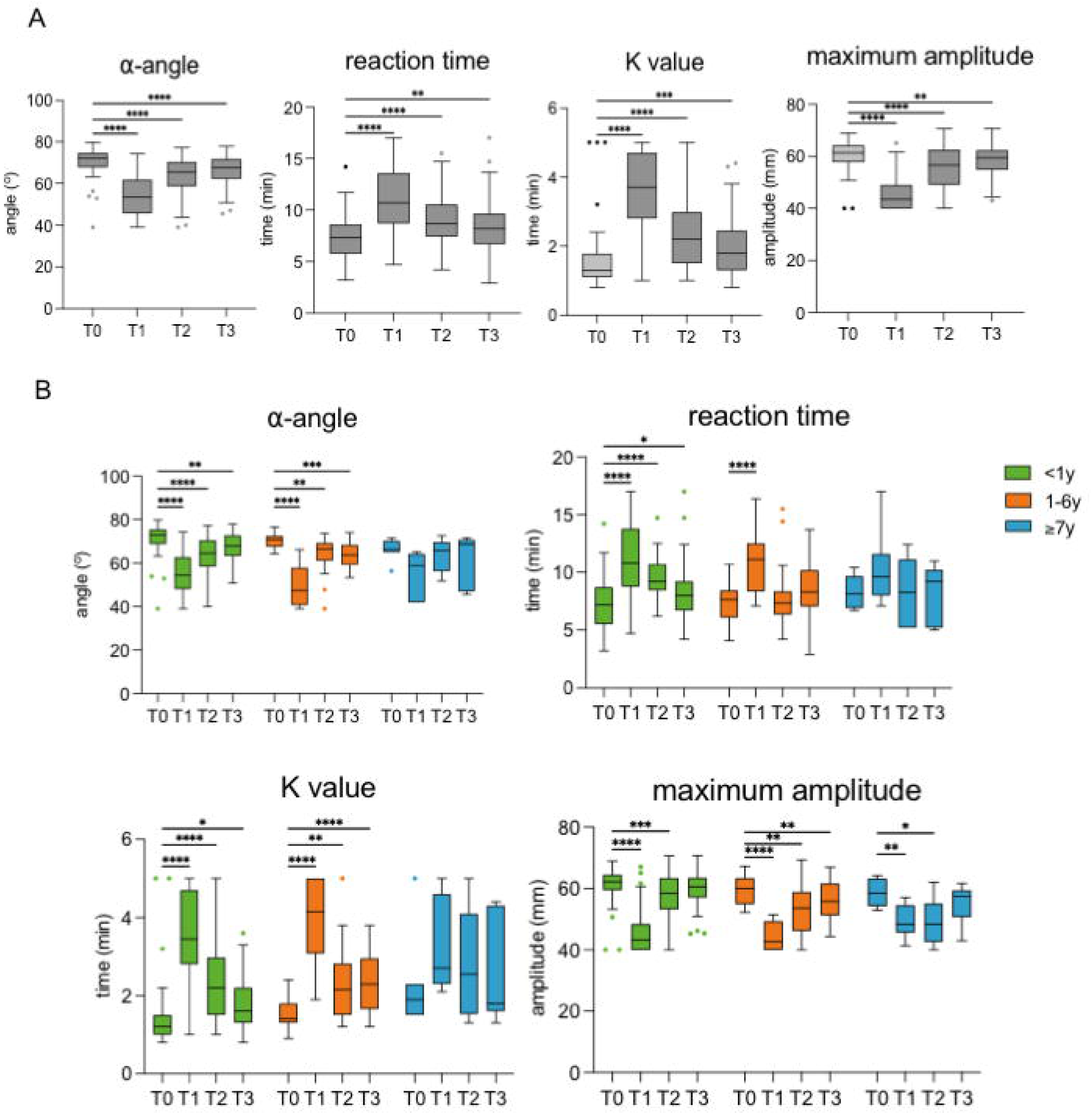
RNA sequencing analysis of neutrophils in neonates and infants at T0 and T2. **(A)** The volcano plot shows upregulated and downregulated genes; the dotted line represents the threshold line of differential gene screening criteria. **(B)** GO biological process enrichment analysis of top 20 significantly up- and down-regulated differentially expressed genes (DEGs). The upper panel showed upregulated DEGs, and the lower panel showed downregulated DEGs. Bar plots represent −log (*p* value) of upregulated and downregulated DEGs. Line and dotted plots represent gene counts of upregulated and downregulated DEGs. **(C)** KEGG pathway enrichment analysis of significant DEGs. Upper panel shows upregulated DEGs and lower panel shows downregulated DEGs. Rich factor refers to the ratio of the number of DEGs to the number of total annotated genes in the pathway. The point color represents −log (*p* value), and the point size represents the number of DEGs mapped to the reference pathway.

### Neutrophils were more transcriptionally active in neonates and infants perioperatively

We also compared neutrophil RNA transcriptomic profiles in neonates and infants against 1-6 years group and ≥7 years group at baseline and ICU admission. Overall, there were 1,643 and 462 common DEGs in neutrophils of patients aged 1-6 years and ≥7 years, respectively, compared to neonates and infants (**Fig. 3A**). By extracting inflammation-related GO terms ^33^, we compared the distribution of gene counts in each term among the three age groups. More gene counts were associated with inflammation-related GO terms in the neonates and infants group compared to the other age groups (**Fig. 3B-C**). Upregulated DEGs showed chemotaxis, migration, adhesion, innate immune response, NFκB signaling pathway, and MAPK cascade (**Fig. 3B**). **Fig. 3D-E** demonstrated GO biological process enrichment analysis of top 20 significantly up- and down-regulated DEGs of patients aged 1-6 years and ≥7 years, respectively. Neutrophils in neonates and infants exhibited amplified transcriptional activity toward inflammatory responses. It is plausible that neonates and infants may be more susceptible to the consequences arising from neutrophil activation after cardiac surgery with CPB.

**Fig. 3.**
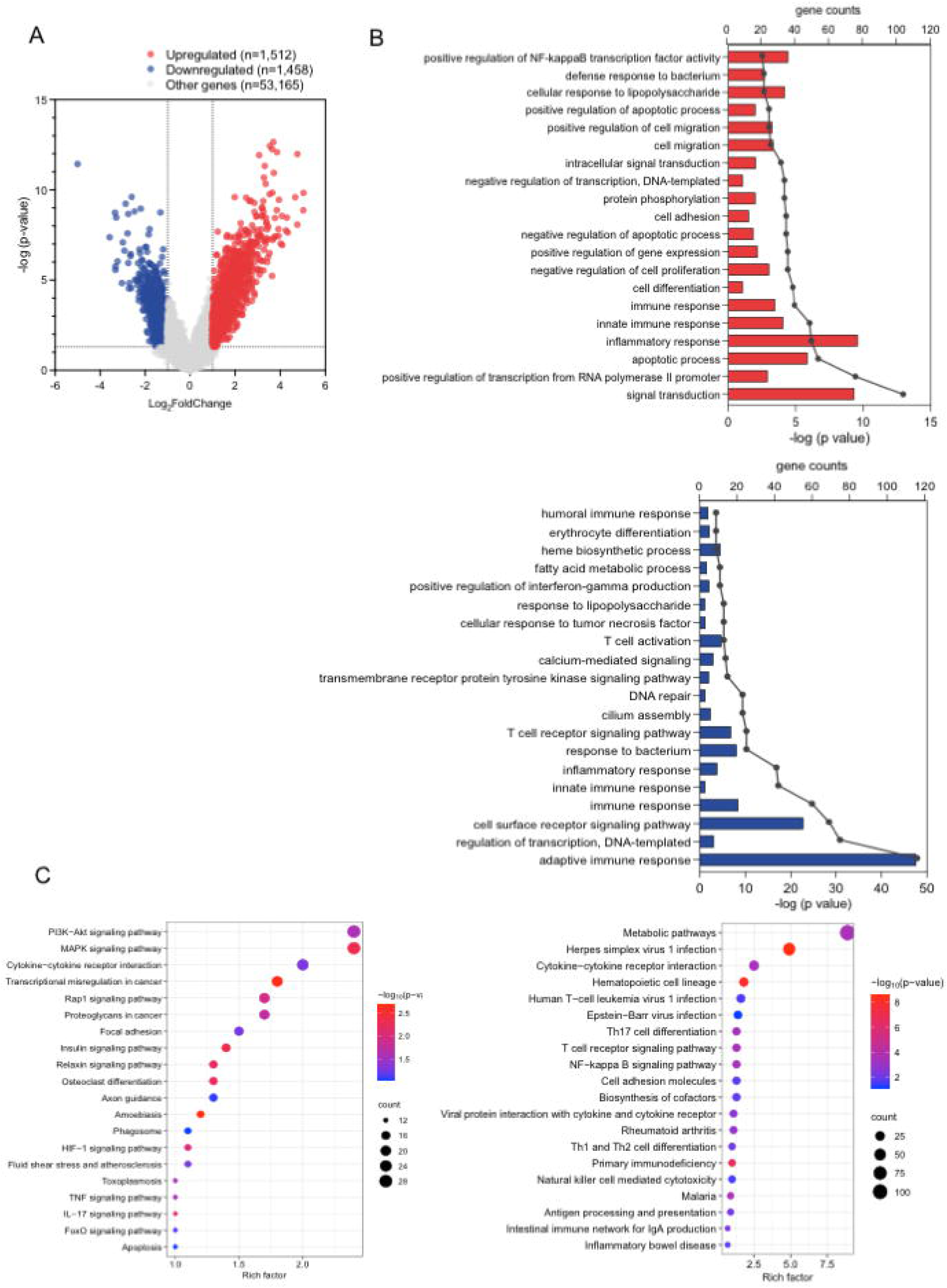
RNA sequencing analysis of neutrophils in neonates and infants at T0 and T2 compared to age 1-6 and ≥7 years. **(A)** Significant DEGs of neutrophils at the ICU admission compared to baseline were determined for each age group. DEGs for neonates and infants were compared to age 1-6 years (upper panel) and age ≥7 years (lower panel), shown on the Venn diagram. Bar graphs illustrate a number of upregulated **(B)** and downregulated **(C)** inflammation-related GO biological processes across different age groups. **(D-E)** GO biological process enrichment analysis of top 20 significantly up- and down-regulated differentially expressed genes (DEGs) of patients aged 1-6 years and ≥7 years, respectively. Right panel showed upregulated DEGs and left panel shows downregulated DEGs. Bar plots represent −log (*p* value) of upregulated and downregulated DEGs. Line and dotted plots represent gene counts of upregulated and downregulated DEGs.

### Neonates and infants undergoing congenital heart surgery developed more multiple postoperative complications

Among a total of 101 patients, 32 patients developed at least one postoperative complication (**Table 2**); 23 patients were neonates/infants (32.9 %) and 9 patients were in the 1-6-year age group (37.5 %). We did not see any complications in patients ≥7 years of age. Neonates and infants were more likely to develop multiple complications (n=12, 17.1%) compared to the 1-6- year age group (n=3, 12.5%). Notably, all cases of thrombosis occurred exclusively in neonates and infants. These fit with the previous report that neonates and infants had highest morbidity and mortality following congenital heart surgery associated with organ injury ^8^. Thrombotic complications and organ dysfunction/failure are responsible for these adverse outcomes ^34^.

**Table 2.** Postoperative characteristics of patients by age groups.

|  | all patients<br>(n=101) | <1y<br>(n=70) | 1-6y<br>(n=24) | ≥7y<br>(n=7) |
| --- | --- | --- | --- | --- |
| Postoperative transfusion<br>(number of patients) |  |  |  |  |
| PRBC | 36 (35.6) | 24 (34.5) | 11 (45.8) | 1 (14.3) |
| Cryoprecipitate | 4 (4.0) | 3 (4.3) | 1 (4.2) | 0 (0) |
| Platelet | 13 (12.9) | 7 (10.0) | 6 (25.0) | 0 (0) |
| FFP | 13 (12.9) | 10 (14.3) | 13 (12.5) | 0 (0) |
| Mechanical ventilation<br>duration (h) | 43.7<br>(18.8,83.1) | 43.6<br>(19.2,71.0) | 48.3<br>(7.4,93.1) | 22.5<br>(20.7,26.6) |
| ICU stay (h) | 92.0<br>(46.0,172.0) | 87.0<br>(46.5,163.5) | 120.0<br>(48.0,190.0) | 65.0<br>(43.0,90.1) |
| Hospital stay (d) | 9.9 (6.0,18.0) | 10.1 (6.0,19.0) | 9.9 (4.9,22.7) | 8.0 (6.8,11.9) |
| Number of complication |  |  |  |  |
| ≥2 | 15 (14.9) | 12 (17.1) | 3 (12.5) | 0 (0) |
| 1 | 17 (16.8) | 11 (15.7) | 6 (25.0) | 0 (0) |
| 0 | 69 (68.3) | 47 (67.1) | 15 (62.5) | 7 (100.0) |
| Type of complication |  |  |  |  |
| respiratory dysfunction | 21 (20.8) | 15 (21.4) | 6 (25.0) | 0 (0) |
| cardiovascular dysfunction | 14 (13.1) | 10 (14.3) | 4 (16.7) | 0 (0) |
| neurologic dysfunction | 3 (3.0) | 1 (1.4) | 2 (8.2) | 0 (0) |
| coagulopathy | 3 (3.0) | 2 (2.9) | 1 (4.2) | 0 (0) |
| renal dysfunction | 1 (1.0) | 1 (1.4) | 0 (0) | 0 (0) |
| hepatic dysfunction | 1 (1.0) | 1 (1.4) | 0 (0) | 0 (0) |
| thrombosis | 4 (4.0) | 4 (5.7) | 0 (0) | 0 (0) |
| PELOD-2 score |  |  |  |  |
| postoperative day 1 | 3.0 (2.0,5.0) | 3.0 (1.5,5.0) | 4.0 (2.0,5.0) | 3.0 (2.0,6.0) |
| postoperative day 2 | 3.0 (0.0,5.0) | 2.0 (0.0,5.0) | 3.0 (0.0,5.0) | 3.0 (2.0,4.0) |
| In-hospital death | 4 (4.0) | 3 (4.3) | 1 (4.2) | 0 (0) |

Data are presented as number (%), median (IQR)
Abbreviations: FFP: fresh frozen plasma; PRBC: packed red blood cell; PELOD-2 score: PEdiatric Logistic Organ Dysfunction (PELOD) score.

### Prolonged CPB time was significantly associated with postoperative complications in neonates/infants

**Table 3** summarized a comparative analysis, examining the association between postoperative complications and perioperative factors in neonates/infants. Patients who developed postoperative complications had significantly lower body weight (p=0.04), higher STAT mortality category (p<0.001), prolonged operative (p=0.001) and CPB times (p=0.001). Upon adjusting for confounding factors, prolonged CPB time emerged as a sole factor independently associated with postoperative complications. Each 30-minute increase in CPB time was associated with a 1.46 times higher odds of developing postoperative complications (95% confidence interval 1.01-2.10, p=0.042). This multivariate analysis demonstrated a good calibration (Hosmer-Lemeshow test p=0.342) and excellent discrimination, with an area under the receiver operating curve of 0.840 (95%CI 0.722-0.958, p<0.001). In addition, we demonstrated significant increases in mechanical ventilation duration (p=0.001), ICU stay (p=0.001), and hospital stay (p=0.001). These findings underscored the role of CPB in the development of postoperative organ dysfunction and thrombotic complications in neonates and infants. We also profiled cytokine levels. Consistent with the outcome profile, injury marker interleukin (IL)-6 was significantly elevated in patients with complications at the ICU admission (**Suppl Fig. 1**). Neutrophil chemoattractant IL-8 was also significantly elevated in the complication group.

**Table 3.** Comparison of patient characteristics by postoperative complication.

|  | Postoperative complication |  | P value |
| --- | --- | --- | --- |
|  | Yes (n=23) | No (n=47) |  |
| Gender: male | 12 (52.2) | 26 (55.3) | 0.80 |
| Age (mo) | 2.5 (0.3,4.0) | 4.0 (0.5,5.9) | 0.08 |
| Body weight (kg) | 4.5 (2.9,5.9) | 5.0 (3.6,6.6) | 0.04* |
| STAT mortality category |  |  | <0.001* |
| 1 | 2 (9.1) | 22 (46.8) |  |
| 2 | 4 (18.2) | 19 (40.4) |  |
| 3 | 8 (36.4) | 2 (4.3) |  |
| 4 | 6 (27.3) | 3 (6.4) |  |
| 5 | 2 (9.1) | 1 (2.1) |  |
| Operative time (min) | 386.0 (310.0,366.0) | 303.0 (279.5,353.6) | 0.001* |
| CPB time (min) | 219.0 (131.0,267.0) | 143.0 (112.0,353.6) | 0.001* |
| Aortic cross-clamp time (min) | 78.0 (65.0,156.0) | 90.0 (74.0,129.0) | 0.79 |
| Circulatory arrest | 6 (26.1) | 6 (12.8) | 0.19 |
| Regional perfusion | 9 (39.1) | 8 (17.0) | 0.07 |
| MUF | 21 (91.3) | 41(87.2) | 0.62 |
| Postoperative transfusion<br>(number of patient) |  |  |  |
| PRBC | 8 (34.8) | 16 (34.8) | 0.95 |
| cryoprecipitate | 1 (4.3) | 2 (4.3) | 0.98 |
| platelet | 4 (17.4) | 3 (6.4) | 0.21 |
| FFP | 6 (26.1) | 4 (8.5) | 0.07 |
| Mechanical ventilation duration (h) | 115.8 (48.7,215.6) | 26.5 (18.0,48.0) | <0.001* |
| ICU stay (h) | 216.0 (93.0,359.0) | 69.0 (45.0,115.0) | <0.001* |
| Hospital stay (d) | 30.8 (12.0,50.1) | 8.9 (5.9,12.6) | <0.001* |

Data are presented as number (%), median (IQR)
Abbreviations: CPB: cardiopulmonary bypass; FFP: fresh frozen plasma; ICU: intensive care unit; MUF: modified ultrafiltration; PRBC: packed red blood cell;

### Neutrophil and platelet counts were comparable in patients with and without postoperative complications

Neutrophils and platelets are two major cell types responsible for thrombosis formation ^23,35-37^. Our statistical analysis did not reveal any significant differences between the two groups, neither in the overall cohort (**Suppl Fig. 2A**) nor in any of the age-specific subgroups (**Suppl Fig. 2B- C**).

### Neonates and infants with postoperative complications exhibited significantly lower MA at T1 and higher MA at T2

Because the coagulation status could affect organ dysfunction, we examined the relationship between complications and TEG values. In the overall cohort, TEG results demonstrated significantly lower R time and MA at T1 in both complication and non-complication groups (p<0.05) (**Suppl Fig. 3A**), followed by some increase in T2 toward their baseline values. In a subgroup analysis, neonates and infants who experienced postoperative complications exhibited a significantly lower MA at the T1 time point (p< 0.05) but a significantly higher MA at the T2 time point (p<0.001) (**Suppl Fig. 3B**), while patients aged 1-6 years did not demonstrate such an association (**Suppl Fig. 3C**). Consistent with this finding, neonates/infants in the complication arm received more platelet, cryoprecipitate, and red blood cell transfusion intraoperatively (**Suppl Fig. 3D**), which may explain their higher MA values (**Suppl. Fig. 3B**) and fibrinogen levels (**Suppl Fig. 3E**) at the ICU admission. Despite the transfusion, they showed higher chest tube output in the postoperative period (**Suppl Fig. 3F**), requiring more transfusion (**Fig. 6G**). Youden-J analysis showed that MA <44 mm at T1 and MA> 56 mm at ICU admission were associated with postoperative complications (**Suppl Table 1**).

### DAMPs were present in plasma following CPB

We examined plasma from neonates and infants using mass spectrometry-based proteomics analysis with the hypothesis that DAMPs would be responsible for complications. We identified an array of DAMP proteins (**Fig. 4A**). The heatmap analysis illustrated distinct temporal patterns of DAMP release after surgery. Histones, the nuclear protein that provides structural support for a chromosome ^38^, exhibited early presence shortly after the reperfusion. Heat shock proteins (HSPs) were highly detectable at ICU admission. Furthermore, amyloids were present on postoperative day 1. The underlying mechanism of these temporal patterns needs future investigation. Along with plasma DAMP profiles, the expression of several DAMP-sensing receptor ^13,39^ was elevated at ICU admission based on neutrophil RNA seq analysis (**Fig. 4B**). We also examined coagulation factors, but did not note clear difference between complication and non-complication groups (**Suppl. Fig 5**).

**Fig. 4.**
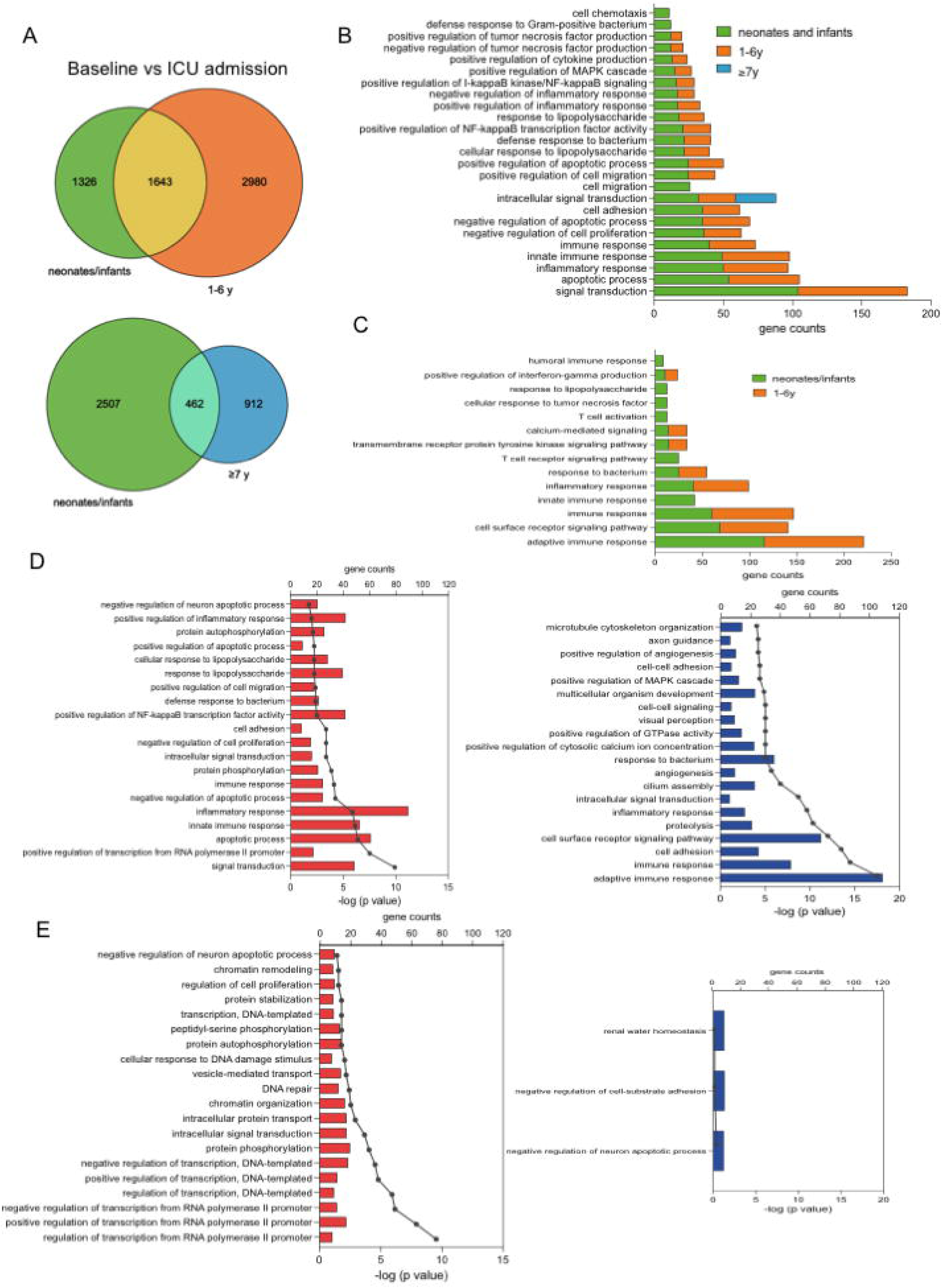
Proteomics-based expression profiles of DAMP molecules in neonates and infants following congenital cardiac surgery with CPB and neutrophil DAMP-related gene expression. **(A)** Heatmap illustrating the temporal changes in the relative expression of DAMPs molecules from T0 to T3, assessed by mass spectrometry analysis of plasma samples. **(B)** Heatmap representing the relative RNA expression of neutrophils compared between T0 and T2, in the overall cohort and subgroup, as determined by RNA sequencing analysis. Bold letters indicate differentially expressed genes (|log2(FoldChange)| >1 and p value <0.05).

### Levels of DAMP molecules were relatively elevated in patients with postoperative complications

To test our hypothesis that DAMPs would be elevated in patients with complications, we used a comparative heatmap analysis (**Fig. 5A**). Several DAMP molecules, including histones (H1, H2- A, H2-B, and H4), HSPs70, HSP90, protein S100A8/A9, serum amyloid proteins, annexin and cathepsin D, exhibited relatively higher expression levels in neonates and infants with postoperative complications compared to baseline levels. We measured plasma histones and S100A8/A9 levels along with plasma HMGB1 levels, whose elevation was described in adult cardiac surgery ^40^. All of them were significantly higher in neonates and infants who developed postoperative complication (**Fig. 5B**). Histones and HMGB1 were already higher at the end of CPB in the complication group.

**Fig. 5.**
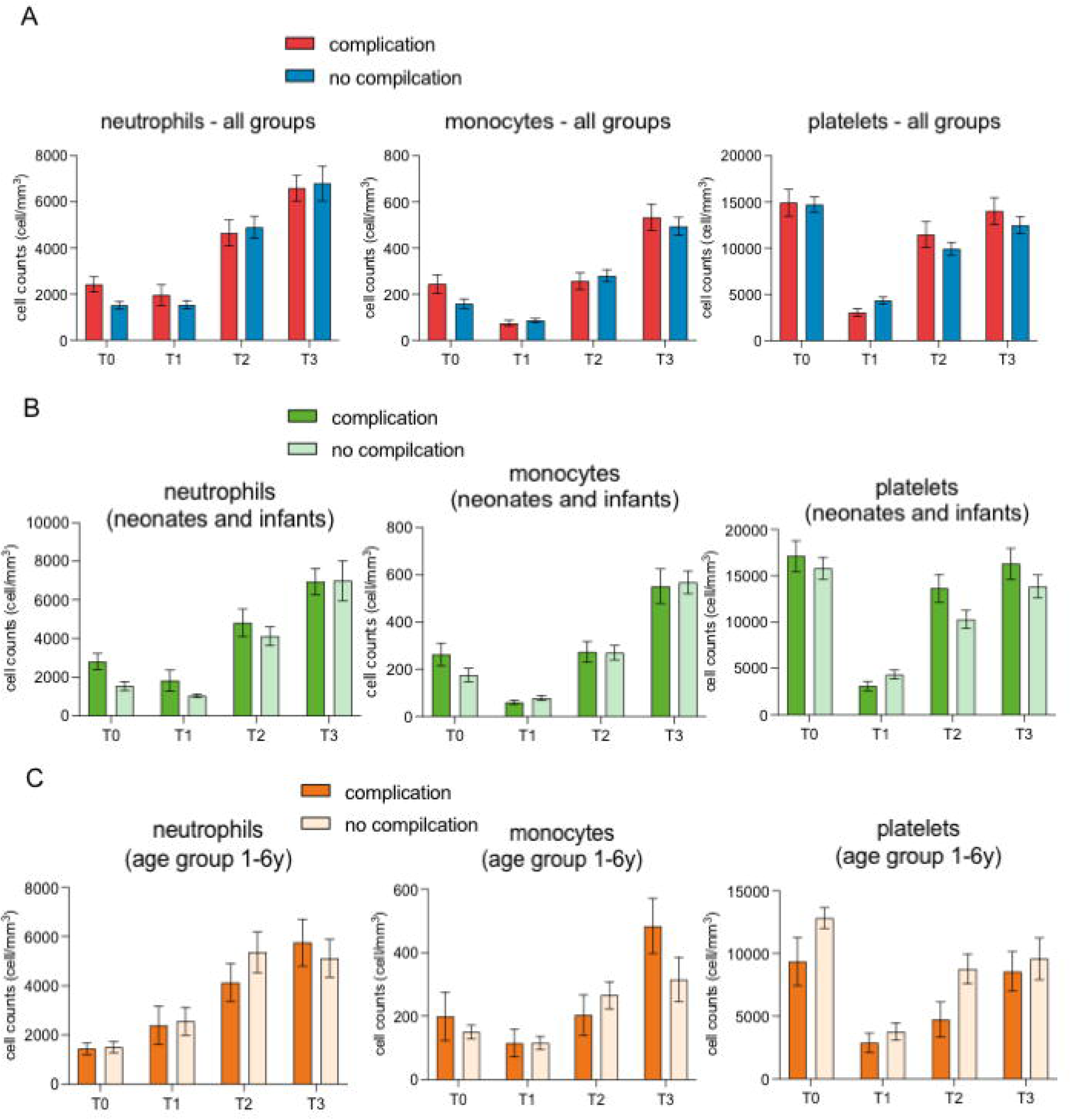
Plasma proteomic analysis of neonates and infants following congenital cardiac surgery with CPB. **(A)** Heatmap illustrating the temporal changes in the relative expression of DAMPs molecules from T0 to T3, comparing neonates and infants with and without postoperative complications. **(B)** Plasma concentration of HMGB1, histones and S100A8/A9 measured by ELISA compared between neonates and infants with and without postoperative complications at T0, T1, T2, and T3. **(C-E)** KEGG pathway enrichment analysis of differentially expressed proteins (log_1.5_(FoldChange) >1) showing upregulated proteins and their associated pathways at T1, T2, and T3 in relation to T0. Rich factor refers to the ratio of the number of differentially expressed proteins to the number of total annotated proteins in the pathway. The point color represents −log (p value), and the point size represents the number of DEGs mapped to the reference pathway. **(F)** Double stranded DNA concentration compared among different age groups. **(G)** Double stranded DNA concentration compared between neonates and infants with and without postoperative complications at T0, T1, T2, and T3. **(H)** MPO-DNA complex levels compared between neonates and infants with and without postoperative complications at T0, T1, T2, and T3. (I) Correlation between MPO-DNA complex levels and histones (or HMGB1). **(B, F, G, H)** Data are presented as mean ± SEM. Statistical analysis was performed using two-way analysis of variance (ANOVA) with multiple comparisons test and *post hoc* Sidak’s correction. *, **, and **** denote p<0.05, 0.01, and 0.0001, respectively. (I) Linear regression analysis was performed. R^2^ and p values were presented.

We performed a KEGG pathway analysis of the upregulated proteins. Our analysis showed that the upregulated proteins were associated with key pathways such as NETs formation and platelet activation during rewarming period **(Fig. 5C)** and upon ICU admission **(Fig. 5D)**, whereas the pathway of complement and coagulation cascade was the most remarkable pathway on the first postoperative day (**Fig. 5E**). Circulating double-stranded deoxyribonucleic acid (dsDNA) has been used as a surrogate of NETs ^41^. dsDNA levels were significantly increased during the rewarming period of CPB and at ICU admission. dsDNA levels were higher in neonates and infants, compared to preschool and school age children (**Fig. 5F**), compatible with our data that neonates and infants had highest complications. Furthermore, dsDNA levels were significantly higher in neonates and infants who developed postoperative complications (**Fig. 5G**). Myeloperoxidase (MPO)-DNA complex level has also been used for NETs measurement ^42,43^. MPO-DNA levels were also significantly higher in the complication group (**Fig. 5H**). Histones and HMGB1 levels were correlated with dsDNA levels and MPO-DNA complex levels (**Fig. 5I**).

### Histones and HMGB1 induced organ injury and thrombosis in mice

As IL-6 was already higher at ICU admission, we focused on histones and HMGB1 whose levels were already elevated intraoperatively. To understand the role of these DAMPs *in vivo*, we conducted intravascular injection of exogenous histones and HMGB1 in mice. Notably, mice injected with histones at a dose of 50 mg/kg died within 5 minutes. Mice injected with histones at a dose of 20 mg/kg or with HMGB1 (0.5 mg/kg) survived for up to 4 hours before euthanasia for subsequent organ histopathological analysis (**Fig. 6A**). Histological examination of the lung in mice injected with histones revealed generalized thrombi formation, accompanied by occasional neutrophil deposition within the thrombi. Injection of HMGB1 also induced thrombus formation in the lung, albeit to a lesser extent compared to histone administration. The lung tissue of mice injected with HMGB1 exhibited noticeable thickening of alveolar septa, neutrophil infiltration within alveolar capillaries, and vascular congestion accompanied by neutrophil recruitment. In the liver, vascular congestion, neutrophil recruitment, and sinusoidal neutrophil infiltration were observed. We also probed NETs formation on the lung. We observed merging of citrullinated histone, myeloperoxidase and DNA, suggesting NETs formation on the lungs in histone (**Fig. 6B**) and HMGB1 injected mice (data now shown). To determine human relevance, we also performed *in vitro* experiments using histone and HMGB1 in human neutrophils. We found that histone and HMGB1 induced NETs (**Fig. 6C**). These results indicate that histones and HMGB1 are major players of NETs formation in pediatric cardiac surgery to cause organ injury. NETs can possibly affect coagulation status through consumption. However, we also noted that histones and HMGB1 lowered platelet counts (**Suppl Fig. 5**). This is in line with lower MA at the CPB rewarming phase in the complication arm and needs further mechanistic investigation.

**Fig. 6.**
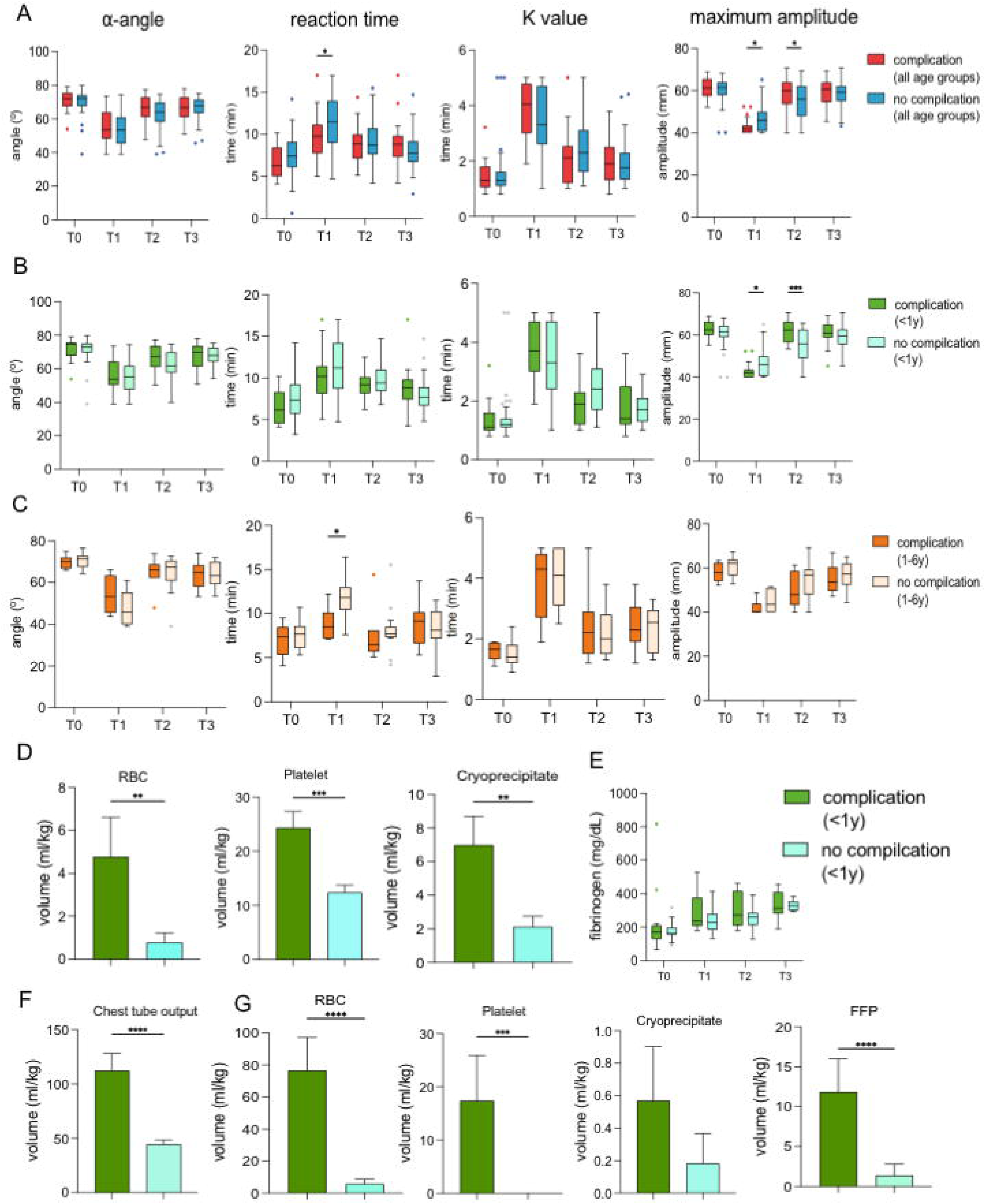
DAMPs induced organ injury, thrombosis, and NET formation in neonates and infants. **(A)** Hematoxylin and eosin (H&E) staining of lungs and liver after intravenous injection of histone and HMGB1. Scale bar (black, 50 µm) was shown in each image; arrows: neutrophils, asterisk: thrombus. **(B)** cit-histone (green), MPO (red) and DAPI (DNA, blue) were co-stained fluorescently in the lung from mice injected with histone. Scale bar (white, 50 µm) was shown in each image. Triple stained part suggested NETs formation. **(C)** *In vitro* NETs experiment stimulated with PMA (50 µM), histone (100 µg/mL) and HMGB1 (50 µg/mL). Arrows indicate NETs.

## Discussion

Here we first delineated the longitudinal immunological responses in pediatric patients undergoing congenital cardiac surgery on CPB. Our data showed a significant increase in neutrophil and monocyte counts across all patients in response to CPB, consistent with the previous studies ^44,45^. We also observed distinct dynamics in neutrophil and monocyte cell number changes based on the age group. Younger children showed more profound, long-lasting neutrophil and monocyte increase in peripheral circulation. Our RNA seq analysis revealed that neonates and infants had most inflammatory DEGs, indicating that their neutrophils might demonstrate exaggerated systemic inflammation. Excessive neutrophil infiltration is associated with tissue damage in a large variety of clinical settings including acute lung injury, myocardial infarction, and cerebrovascular disease^46-49^, which could be in support of higher morbidities and mortalities in neonates and infants following cardiac surgery.

As in the Kids’ Inpatient Database (KID), neonates and infants undergoing cardiac surgery on CPB had in-hospital mortality rate of 6.9%, associated with thrombosis and organ injury, while the rates of age 1-5, 6-12, and 13-17 years were 1.28%, 0.67%, and 0.83%^8^. As in line with this report, our data showed neonates and infants had highest rate of multiple complications postoperatively with an overall mortality rate of 4%. Specifically, the mortality rates for neonates/infants and the age group of 1-6 years were 4.3% and 4.2% respectively. DAMPs are the instigators of neutrophil activation^26^ and contribute to the pathogenesis of systemic inflammation and thrombogenesis, leading to organ dysfunction and immunothrombosis-associated microcirculatory abnormality^50,51^. DAMPs are typically released from damaged tissue to its local environment. During CPB, the release of DAMPs can be amplified due to a period of organ ischemia and subsequent cellular death. Upon reperfusion, these localized DAMP molecules may be disseminated systemically. Several DAMP-sensing PRRs were significantly differentially expressed from circulating neutrophil RNA after CPB exposure. In particular, TLR2 and TLR4 are responsible for major DAMPs molecules, including histones, HMGB1, HSPs, and S100 proteins. DAMP-TLR interaction initiates intracellular signaling cascades, resulting in the production of pro-inflammatory cytokines, chemokines, and the recruitment and activation of additional immune cells. We also identified other endogenous molecules including components of the extracellular matrix (ECM). Although these molecules have shown potential as diagnostic and prognostic markers for inflammatory diseases, their roles remain to be determined^52,53^.

DAMPs can affect multiple cells including neutrophils and platelets, both of which express TLR2 and TLR4. Upon exposure to DAMPs, neutrophils undergo activation^54^, resulting in the production of reactive oxygen species (ROS), release of granule contents, and release of NETs^55-57^. As dsDNA levels were highest in neonates and infants, NETs formation plays a critical role in postoperative complications. Furthermore, in neonates and infants with complications, their dsDNA levels were significantly higher, suggesting the occurrence of NETs. In accordance with these findings, we showed that histones and HMGB1 induced NETs *in vitro* and *in vivo*. NETs primarily originate from the nucleus, leading to a high enrichment of core histones. They also contain abundant granule proteins (neutrophil elastase, cathepsin G, and proteinase), as well as cytosolic proteins such as myeloperoxidase (MPO) and S100 proteins^58^. A growing body of evidence suggested that NETs contributed to arterial and venous thromboses as well as disseminated intravascular coagulation (DIC)^59-62^. Mechanistically, after NET formation, DNA released intravascularly is both procoagulant and cytotoxic. Proteins that bind to DNA including histones and HMGB1, are also procoagulant^63^. NETs provides a scaffold for platelets and erythrocytes to adhere, inducing platelet aggregation and permitting fibrin accumulation contributing to DIC pathogenesis^64^. The intravenous administration of exogenous histones and HMBG into mice also induced organ injury. Histological examination of the lung and liver after injection revealed acute inflammation characterized by vascular congestion, alveolar wall thickening, and neutrophil recruitment, consistent with the early process of multiple organ injury. Intravascular thrombi were observed in the lungs following administration of exogenous DAMPs. This result was consistent with the previous literature reporting that histones and HMGB1 induced thrombosis in both human and animal models ^15,65,66^. Tissue/organ injury as a result can induce further release of DAMPs. Platelets also contribute to NETs. Activated neutrophils also interact with platelets through receptor such as P-selectin, GPIIb/IIIa, and GPIbl1^22^, leading to the formation of platelet-neutrophil aggregates. These aggregates serve as a platform for the induction of further neutrophil activation and priming which will initiate the vicious cycle. Interestingly, histones and HMGB1 induced a reduction in platelet counts irrelevant of NETs. Consistent with our findings, histone-induced thrombocytopenia was previously reported ^67^. This is consistent with lower MA at the end of CPB in neonates and infants with complications. This finding is clinically important, because patients with complication received more transfusion not only intraoperatively but also postoperatively. Histones can also activate platelets through TLR2 and TRL4, initiate the coagulation cascade to convert prothrombin to thrombin, and stimulate fibrin formation by interacting with red blood cells ^68^. This may lead into the consumption of clotting factors. Based on our data and consideration above, we proposed the scheme presented in **Fig. 7**.

**Fig. 7.**
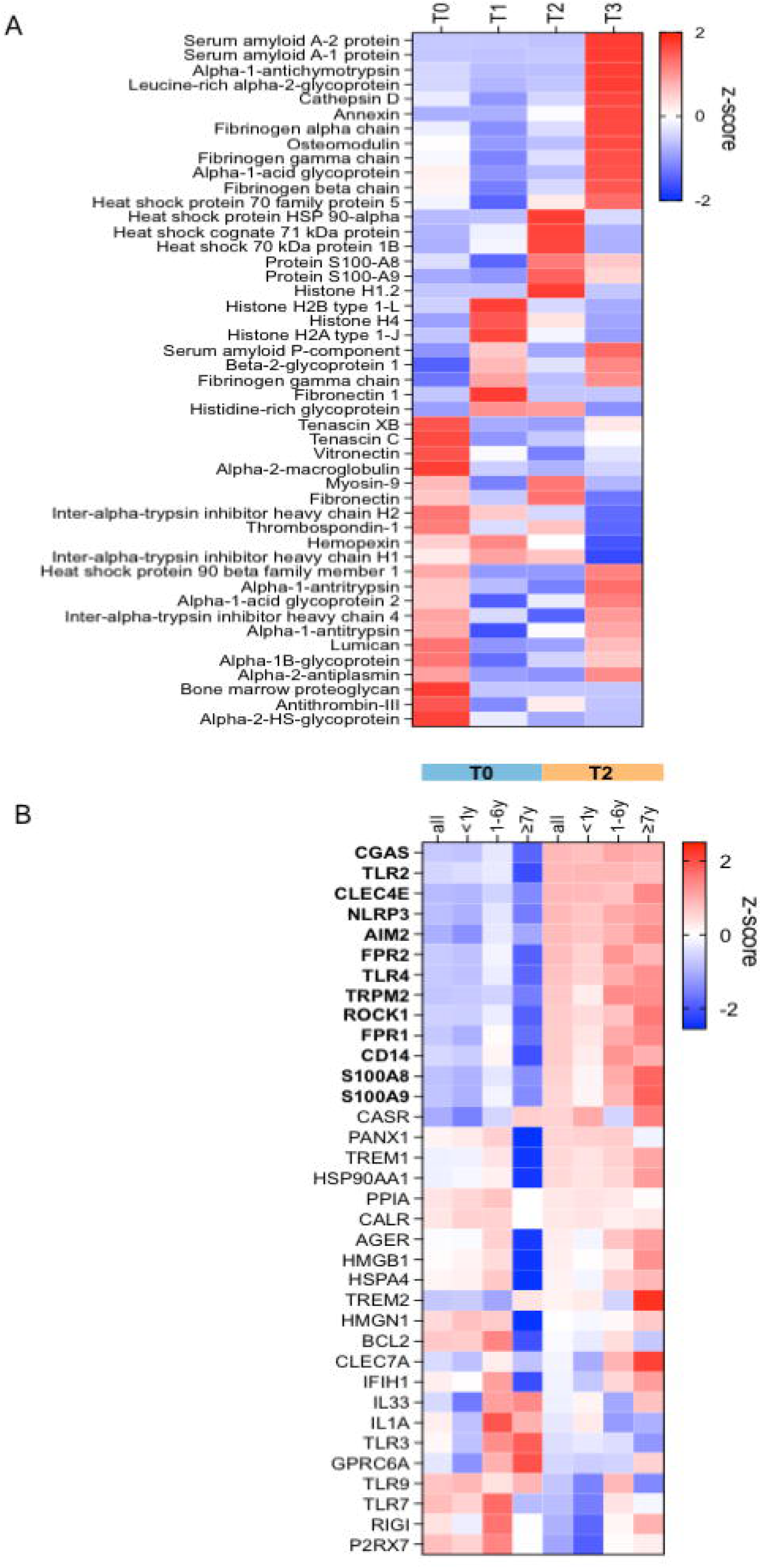
Proposed mechanism of tissue/organ injury and transfusion requirement in patients with complications. Patients with complications demonstrated higher DAMP levels (including histones, HMGB1), which induced platelet consumption and neutrophil activation. Activated neutrophils can induce NETs, which further cause tissue/organ injury due to thrombosis, leading to more DAMP production. To compensate for platelet consumption, more transfusion is required.

**Figure 8.**
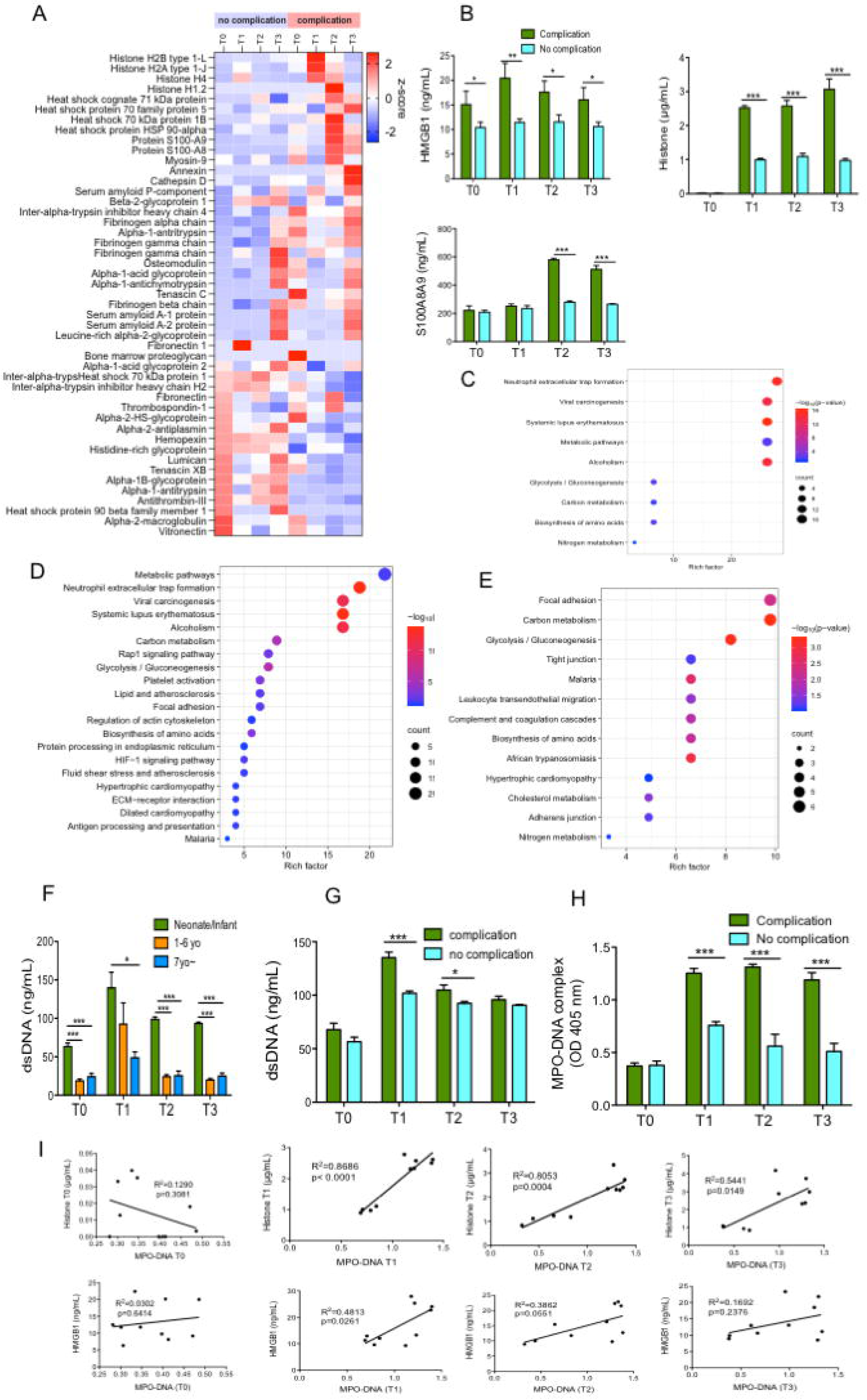

**Figure 9.**
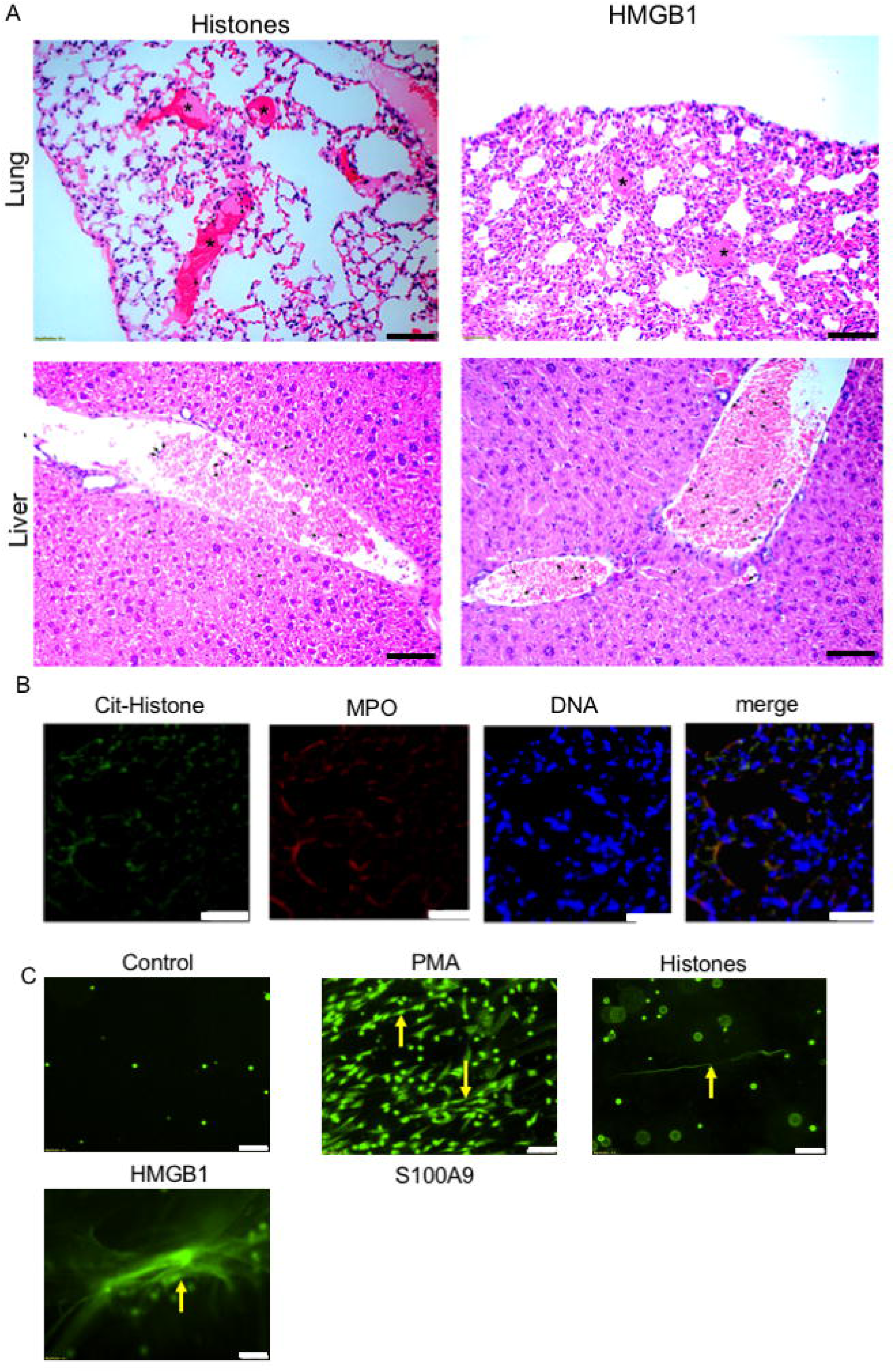

**Figure 10.**
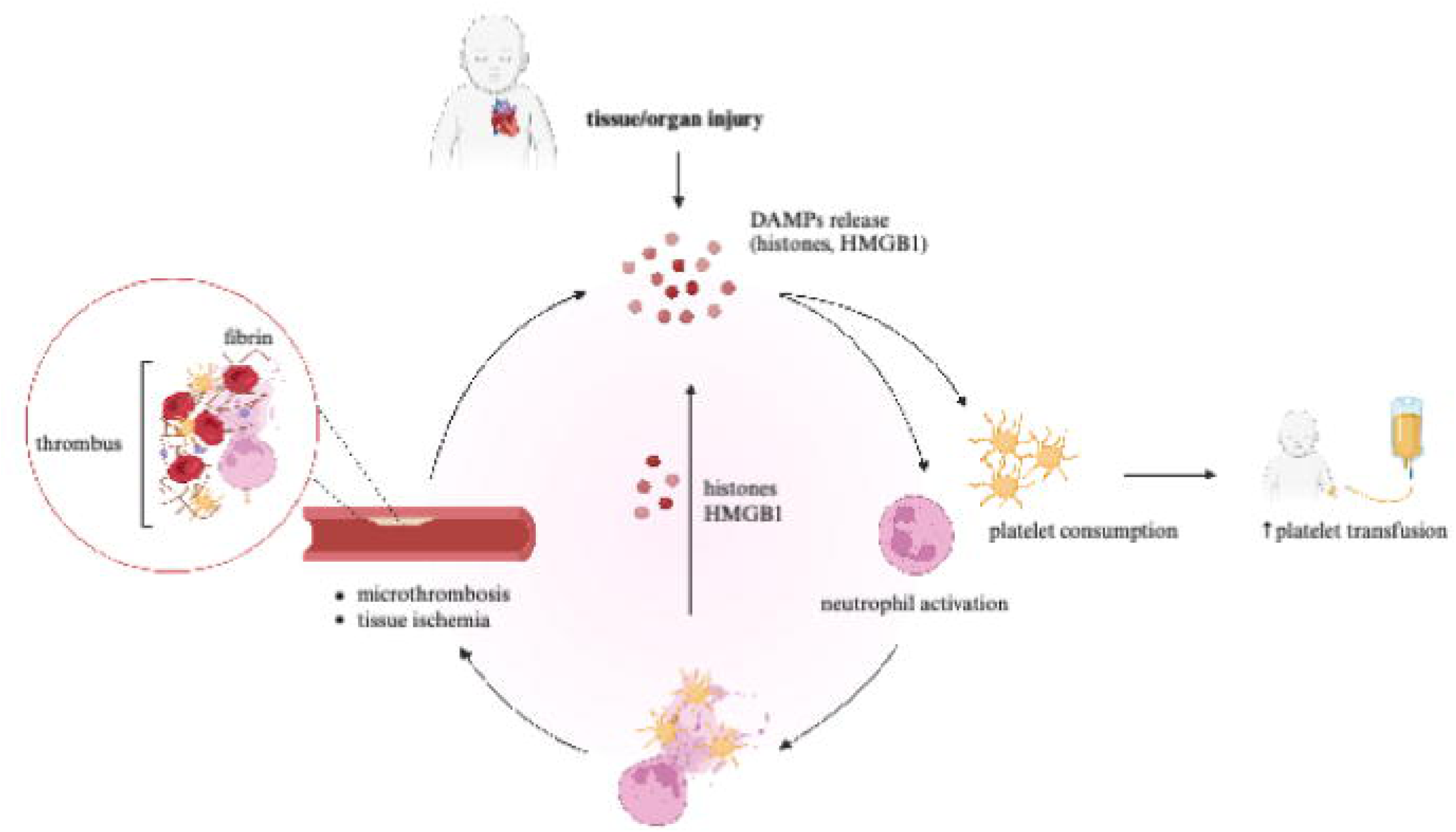

The association between DAMPs, NETs, and postoperative organ dysfunction and thrombosis is of significant interest. Certain DAMPs have a capacity to trigger remote thrombotic and inflammatory insults in distant organs from the primary injury, serving as both diagnostic biomarkers and prospective therapeutic targets for preventing organ injury and thrombosis^49,65,69^. Notably, histone administration resulted in the most prominent pulmonary thrombi, which correlated with early mortality in the mice. The distribution of circulating DAMPs injected intravenously could be varied among different organs, with the lungs showing higher susceptibility compared to the liver. This observation is consistent with our clinical data, where respiratory dysfunction was the most prevalent complication in the cohort. The organ-specific effects of DAMPs might be influenced by the distinctive physiological characteristics of capillaries within each organ, which play a significant role in determining susceptibility to tissue injury^15^. Furthermore, the differences in the severity of inflammation and thrombus formation following the infection of each DAMP molecules might be attributed to dose-dependent and time-dependent effects, as previously demonstrated with histones^15^. Therefore, further investigations are warranted to assess the effects of DAMPs at different doses and their temporal dynamics. It is also important to note that not all neutrophils can undergo NETs formation under the same stimulation because the activation of human neutrophil results in only 60% NETs formation^70^, thus, not all patients will manifest organ dysfunction or thrombosis even they have similar degrees of neutrophil activation. Environmental signals and intrinsic characteristics or the combination of both, may alter the propensity of certain neutrophil subpopulation to release NETs in response to stimuli^49,71^.

While extensive research has focused on the pathological mechanism of individual DAMP molecules through *in vitro* and *in vivo* experiments, the comprehensive characterization of DAMPs in clinical scenarios remains limited. In this sense, our study is unique by identifying multiple types of DAMPs that are concurrently released and interact with various PRRs. However, the intricate functional interplay and the possible synergistic, additive, or competitive effects of these DAMPs remain incompletely elucidated. We also recognize the heterogeneity of patients enrolled in our cohort, which is often inherent to cardiac surgical cases and may potentially influence the observed outcomes.

In conclusion, our study provides compelling evidence for the involvement of DAMPs and NETs in the development of postoperative organ dysfunction and thrombosis in neonates and infants undergoing congenital cardiac surgery with CPB. Our findings shed light on the pathogenesis of postoperative complications and highlight the potential targeting DAMPs and NETs as preventive and therapeutic strategies to mitigate organ dysfunction and thrombotic events following congenital cardiac surgery with CPB.

## Data availability

All the data are presented in this manuscript.

## Acknowledgement

This is supported by NICHD R21HD109119 (K.Y.)

## Notes

Conflict of Interest None

### Competing Interest Statement

The authors have declared no competing interest.

