## Supplementary material for "Neutrophil extracellular traps formation is associated with postoperative complications in neonates and infants undergoing congenital cardiac surgery": Supplmental Figures … table

Short title: NETs and congenital cardiac surgery

Wiriya Maisat^1,2,3,4^, Lifei Hou^1,2,3^, Sumiti Sandhu^5^, Yi-Cheng Sin^6^, Samuel Kim^1^, Hanna Van Pelt^1^, Yue Chen^6^, Sirisha Emani^7^, Sek Won Kong^5,8^, Sitram Emani^7^, Juan Ibla^1,2^, Koichi Yuki^1,2,3,9^

^1^Department of Anesthesiology, Critical Care and Pain Medicine, Boston Children’s Hospital, Boston, MA, USA

^2^Department of Anaesthesia, Harvard Medical School, Boston, MA, USA

^3^Department of Immunology, Harvard Medical School, Boston, MA, USA

^4^Department of Anesthesiology, Faculty of Medicine Siriraj Hospital, Mahidol University, Bangkok, Thailand

^5^Computational Health Informatics Program, Boston Children’s Hospital, Boston, MA, USA

^6^Department of Biochemistry, Molecular Biology and Biophysics, University of Minnesota, Minneapolis, MN, USA

^7^Department of Cardiac Surgery, Boston Children’s Hospital, Boston, MA, USA

^8^Department of Pediatrics, Harvard Medical School, Boston, MA, USA

^9^Broad Institute of Harvard and MIT, Cambridge, MA, USA

**Supplemental methods**

*Patient selection and perioperative course*

We included all eligible patients scheduled for congenital cardiac surgery with CPB. Exclusion criteria were patients who did not necessitate CPB, presented with an active infection, received chronic steroid therapy, had immunodeficiency, HIV infection, or a history of malignancy. Patients were included from May 31, 2022, to February 22, 2023.

All patients included in the study underwent general anesthesia with endotracheal intubation and arterial and central venous line placement. Subsequently, after surgical dissection, patients were heparinized and cannulated with aortic and venous cannulas for CPB. The CPB circuit was primed with one unit of packed red blood cells (pRBC) and one unit of fresh frozen plasma (FFP), maintaining a target hematocrit level greater than 30% per our institutional standard protocol. The use of circulatory arrest or regional perfusion, temperature management, and modified ultrafiltration (MUF) was determined on a case-by-case basis. In cases of non-surgical microvascular bleeding, platelets, and cryoprecipitate were administered. Neonates and infants typically did not receive FFP transfusion after CPB in our institution. Following surgery, patients were either extubated or kept intubated based on the discretion of the attending anesthesiologist, and they were subsequently transferred to the intensive care unit (ICU) for postoperative care.

*Clinical data collection*

Demographic information, comorbidities, diagnosis, procedures, medications, laboratory values, CPB details, type and volume of blood products administered, postoperative complications, respiratory support, vital signs, as well as hospital and ICU lengths of stay were extracted from the electronic medical record for the analysis of clinical data. Society of Thoracic Surgeons - European Association for Cardio-Thoracic Surgery Congenital Heart Surgery Mortality Categories (STAT Mortality Categories) was used to estimate the risk of morbidity associated with operations

Postoperative complications included organ dysfunction/failure and/or thrombosis. Due to the absence of a standardized definition for organ dysfunction following congenital cardiac surgery, we adopted previously published criteria established by others^1,2^. Organ dysfunctions included 1) cardiovascular dysfunction (low cardiac output syndrome, reliance on a vasoactive drug to maintain blood pressure, or two of the followings; metabolic acidosis, elevated arterial lactate, oliguria, or prolonged capillary refill), 2) respiratory dysfunction (arterial oxygen tension/fraction of inspired oxygen (P_a_O_2_/F_i_O_2_) < 300, arterial carbon dioxide tension (P_a_CO_2_) > 65 torr or 20 mmHg over baseline P_a_CO_2_, need for > 50% F_i_O_2_ to maintain oxygen saturation ≥ 92%, or need for non-elective mechanical ventilation, prolonged mechanical ventilation ≥ 5 d, 3) renal dysfunction (the presence of creatinine of > 114 μmol/L, urine output of < 1 mL/kg/hr despite diuretic administration, or the requirement for ultrafiltration or hemodialysis 4) coagulopathy or bleeding complication requiring chest exploration for bleeding or removal of clots, intracranial hemorrhage, prothrombin or partial thromboplastin time was three times normal, or > 30 mL/kg of blood products were infused during a 24-hr period, 5) central nervous system dysfunction ( development of a new intracranial infarct or hemorrhage, evidence of hypoxic-ischemic injury by clinical examination or computed tomography of the head, or brain death), 6) hepatic dysfunction (bilirubin concentration of > 2 mg/dL and/or increase of hepatic cellular enzymes two or more times normal). Thrombosis was defined as the presence of any vascular thrombosis detected using ultrasound diagnostic imaging.

*Blood sample collection*

Blood samples were collected from patients through existing central venous catheters. The blood was obtained at four-time points; 1) after anesthesia induction (T0 - baseline), 2) during the rewarming period (T1), 3) upon admission to the ICU (T2), and 4) on postoperative day 1 (T3). The volume of blood required for each collection time point was determined in accordance with the guidelines outlined in the "Protection of Human Subjects" document, with a total volume of 1 mL. TEG analysis was performed on a portion of the blood at the coagulation laboratory, while the remaining blood was promptly transported to the research laboratory in a heparinized tube for further analysis.

*Thromboelastography*

Citrated whole blood samples were collected at four time points (T0, T1, T2, and T3) for TEG. TEG analysis was performed on all samples using TEG^®^500 in a Clinical Laboratory Improvement Amendments (CLIA) certified laboratory. R time in min, k time in min, alpha (⍺) angle (degrees), and MA in mm were recorded for each sample.

*Neutrophil, monocyte, and platelet counts*

A total of 10 μL of whole blood collected at four time points (T0, T1, T2, and T3) was incubated with Fc receptor blocking solution and subsequently stained with fluorescent-conjugated antibodies against CD15-APC, CD14-PE, and CD41-PerC (Biolegend Inc., San Diego, CA, USA). Following a 30-min incubation, the stained samples were treated with 10% FACS™ Lysing Solution (BD Bioscience, San Jose, CA, USA). The cells were then suspended in phosphate-buffered saline (PBS) and subjected to flow cytometry analysis using a BD Accuri C6 flow cytometer (BD Biosciences, Franklin Lakes, NJ, USA). Neutrophils, monocytes, and platelets were identified based on their expression of CD15, CD14, and CD41, respectively. A total of 100 μL of each sample was collected for each analysis.

*RNA sequencing analysis of neutrophil mRNA*

Neutrophils were isolated from whole blood samples collected at T0 and T2 through density gradient separation with Polymorphprep™ according to the manufacturer's protocol (AXIS-SHIELD PoC AS, Oslo, Norway). Briefly, fresh citrated whole blood was layered over an equal volume of Polymorphprep™ (Proteogenix, Miami, FL, USA) and centrifuged at 500 xg for 30 min. The neutrophil fraction was collected and subjected to RNA extraction using Trizol reagent (Life Technologies, Carlsbad, CA, USA) following the manufacturer’s protocol.

RNA sequencing was performed using Illumina TruSeq Stranded Total RNA sample preparation, followed by next-generation sequencing (NGS) using Illumina NovaSeq to generate 50 million 150bp paired-end reads per sample. Library preparation and NGS were performed at the Harvard Institute of Medicine. This sequencing depth provides abundant sensitivity for all expressed transcripts, both protein-coding and regulatory (microRNA and long non-coding RNA), including exon-exon junction reads to detect differential usage of splicing variants. The analysis proceeded using a standard bioinformatics pipeline developed for profiling transcript-level abundance in bulk RNA-seq. The quality of the raw reads was evaluated using FastQC (v0.11.9). Any detected Nextera transposase adapter contaminations within the reads was trimmed using TrimGalore (v0.6.10). The reads were then aligned to the human reference genome GRCh38 using the 2-PASS STAR algorithm (v2.7.10b). HTseq-count (v2.0.2) was employed to generate raw counts for each transcript. Inter-individual differences were found to be significantly more pronounced than the differences between pre and post-surgery states. To account for this subject-level variability, a differential expression analysis was conducted on the log2-normalized expression data using a mixed-effect model with the lmerSeq R package. This model compared post-surgery states to pre-surgery ones. The Bonferroni correction was applied to adjust p-values for multiple comparisons. Genes that exhibited a false-discovery rate less than 0.05 and an absolute fold change greater than 1.5 were considered significant. GO Enrichment analysis was performed to demonstrate the function of significant DEGs in Biological Process, and KEGG pathway were examined using DAVID online analysis tool.

*Inflammatory Cytokine measurement*

Levels of serum cytokines TNF-α, IL-1β, IL-2, IL-4, IL-6, IL-8, IL-10, IL-12p70 and IL-13 were measured by using human Th1/Th2 (Meso Scale Discovery; Gaithersburg, Maryland) per the company protocol.

*Proteomics sample preparation*

Plasma solution was diluted two folds with urea lysis buffer (9 M urea, 50 mM ammonium bicarbonate with cOmplete™ Protease Inhibitor Cocktail (Roche, Penzberg, Germany)) and incubated at room temperature with rotation for 10 min. Then, each sample was centrifuged at 21,000 x g for 10 min. The proteins in the supernatant was reduced and alkylated with 10 mM Tris(2-carboxyethyl)phosphine hydrochloride (TCEP-HCl) (Pierce, Appleton, WI) and 10 mM iodoacetamide (VWR, Radnor, PA) for 30 min in dark, followed by quenching with 20 mM cysteine for 30 min. Finally, each sample was diluted to 1.5 M urea with water and digested by trypsin at an enzyme-to-substrate ratio of 1:60 (w/w) (Promega, Madison, WI) at 37 °C overnight. A second round of trypsin digestion was performed under the same condition the next day for 2 hrs.

*Nano-HPLC-MS/MS analysis*

Digested peptides were desalted with C18 Stage tips similarly as previously described ^74^ and analyzed in the Center for Metabolomics and Proteomics (CMSP) at the University of Minnesota. Briefly, desalted tryptic peptides were reconstituted in HPLC sample buffer (97.99:2:0.01, water:acetonitrile (can):formic acid (FA)) and loaded onto a self-packed reversed-phase capillary HPLC column (40 cm length x 100 µm internal diameter, ReproSil-Pur Basic C18aq, 1.9 µm particle size, 120 Å pore size (Dr. Maisch GmbH, Ammerbuch, Germany)). Peptides were analyzed on a nano-HPLC-MS system with UltiMate™ 3000 RSLCnano System and Orbitrap Fusion mass spectrometer (ThermoFisher, Waltham, MA). Peptides were eluted over a linear gradient with the following profile: 5% B solvent from 0 – 2 minutes, 8% B at 2.5 minutes, 21% B at 6 minutes, 35% B at 90 minutes and 90% B at 92 minutes with HPLC buffer A (0.1% formic acid in water, v/v) and HPLC buffer B (0.1% formic acid in acetonitrile, v/v). The flowrate profile was 400 nl/min from 0 – 2 minutes, 315 nl/minute from 2.5 – 90 minutes and 400 nl/minute from 90 – 92 minutes. The Orbitrap Fusion mass spectrometer (ThermoFisher, Waltham, MA) was operated with a full scan range of 380–1580 m/z at 120,000 resolutions (at 200 m/z) and HCD MS/MS analysis acquiring the top 12 precursor ions in data-dependent acquisition (charge states from 2 to 6) with an isolation window of 1.6 m/z and fragmentation energy at 35% in the linear ion trap.

*Proteomics data analysis*

Mass spectrometry data and quantitative analysis was performed with MaxQuant software (version 1.5.3.12)^3^. Data was searched against the human proteome database (UP000005640.fasta) concatenated with a common contaminant database from Maxquant and filtered at the 1% false discovery rate (FDR) for both peptide and protein identifications through the reverse target-decoy strategy. A maximum of 1 missing cleavage was allowed for trypsin as the proteolytic enzyme for database searching. The maximum number of modifications per peptide was set at 3. Carbamidomethylation on cysteine was set as a fixed modification, and protein N-termini acetylation and methionine oxidation were set as variable modifications. Label-free quantification analysis (LFQ) was applied for the relative quantification of protein abundance^4^.

*HMGB1, histones and S100A8/A9 assay* *using enzyme-linked immunosorbent assay (ELISA)*

Plasma HMGB1 levels was evaluated with human HMGB1/HMG-1 colorimetric ELISA kit (Novus #NBP2-62766) with some modifications from the manufacturer’s instructions. The absorbance was measured in a plate reader at a wavelength of 450 nm. A calibration curve was generated from OD values of the standards at each concentration to calculate HMGB1 concentrations. Similarly, histones and S100A8/A9 levels were measured by ELISA kits per company instructions (Abcam and R&D systems).

*double-stranded DNA assay*

Serum dsDNA levels were quantified using a fluorescent nucleic acid stain, Quant-iT PicoGreen (Invitrogen), according to the manufactu’er's protocol. Blood samples were prepared based on previous study^5^ to detect an estimated plasma DNA range from 50-200 ng/mL. Fluorescence was excited at 480 nm, and the intensity of emission was detected at 520 nm. Standard curves were generated for the calculation of dsDNA concentrations.

*Effects of histones and HMGB1on organ injury and thrombosis in mice model*

We used a murine model to evaluate the effects of histones and HMGB1on organ injury and thrombosis. Wild type mice on the C57BL6 background were obtained from the Jackson laboratory (Bar Harbor, Maine). They were housed under specific pathogen-free conditions, with 12-h light and dark cycles. All animal protocols were approved by the Institutional Animal Care and Use Committee (IACUC) at the Boston Children’s Hospital. All the experimental procedures complied with the Animal Research Repositing *In Vivo* Experiments guideline^78^. Mice were injected with intravenous histone 20 and 50 mg/kg (Anaspec, Fremont, CA) and HMGB1 50 mg/kg (SinoBiological, #50923-M01H). Experimental animals were sacrificed after 4 hours of the injection. For histopathological analysis, lungs and livers were harvested and fixed in 10% formalin for 48 h. Tissue samples were dehydrated and embedded in paraffin. The embedded samples were cut into thin sections using a microtome, placed the section on microscope slides, stained with hematoxylin and eosin (H&E) solution, and visualized with light microscopy (Olympus IX81).

*Immunofluorescent staining*

Slides were permeabilized with 0.2% Triton-X for 30 min, blocked with 2% normal donkey serum in PBS / 0.2% Triton-X for 1h at room temperature, stained with 1 μg/ml of primary rabbit anti- Histone H3 (ab5103) and Goat-anti-human/mouse MPO (AF3667, R&D system) overnight at 4 degree, washed with PBS-0.1% Tween-20 and incubated for 1h with secondary antibodies NL557 Donkey-anti-Goat IgG and NL493 Donkey-anti-rabbit IgG (both R & D system) at room temperature. The slides were washed with PBS-0.1% Tween-20 and mounted with DAPI-labeled mounting medium and evaluated by fluorescent microscopy.

*In vitro NETs assay*

Human neutrophil was purified from healthy donor-derived peripheral blood through MACSxpress whole blood neutrophil isolation kit according to product instruction (Miltenyi Biotech). Purity was checked to be >90% by flow cytometry. Neutrophil (1 ⋅ 10^5^ in 200 μl) in RPMI-1640 with 10 mM HEPES, 50 µM 2-ME, and 2% FCS (heated at 70°C for 30 min to inactivate the DNase) were allowed to adhere to eight-well glass chamber slides (Thermo Fisher Scientific, Waltham, MA) for 1 hour at 37°C and then were stimulated with various simulators including medium control, phorbol 12-myristate 13-acetate (PMA, 50 nM as a positive control), histone (100 µg/mL) and HMGB1 (50 µg/mL) for 4 hours. At the end of culture, Sytox green (0.1 ⎧M, final concentration) was added and NETs were imaged with a fluorescence microscope, Olympus IX81 Motorized Inverted research microscope system.

*Statistical analysis*

The sample size was determined based on an estimated incidence rate of thrombosis as the outcome variable, with a calculated rate of 9%. To achieve a desired area under the curve (AUC) of 0.75, a minimum of 99 patients were required. For the analysis of clinical data, Stata version 17.0 software (StataCorp, College Station, TX, USA) was used. Categorical variables were presented as numbers and percentages, while continuous variables were summarized as means and standard deviations for normally distributed variables, or medians and interquartile ranges for variables with skewed distributions. The normality of variables was assessed using the Shapiro-Wilk normality test. We compared the proportions of categorical variables with chi-square test, Fisher exact test, or Kruskal-Wallis test as appropriate. The distribution of continuous variables was assessed using Student’s *t* test or Mann-Whitney U test as appropriate. Statistical significance was defined as p < 0.05. All variables with a p value of < 0.05 from univariate analysis were entered into multivariate logistic regression analysis, which was determined using logistic regression with a backward variable selection procedure.

Laboratory data were analyzed as provided in the corresponding figure legends. Statistical significance was defined as p < 0.05. All statistical analyses were performed using Prism 6 software (GraphPad Software, La Jolla, CA, USA).

**Supplemental Figures**

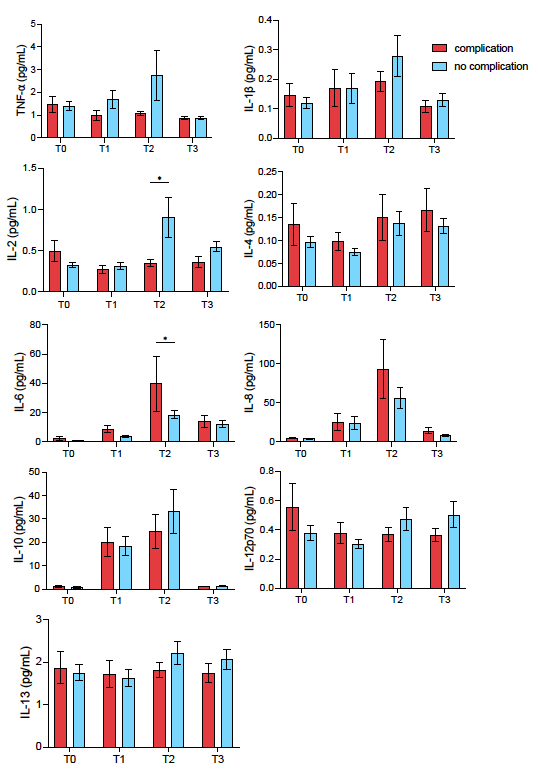

**Suppl Fig. 1. Cytokine measurement in the perioperative period.** TNF-a, IL-1b, IL-2, IL-4, IL-6, IL-8, IL-10, IL-12p70, and IL-13 levels were measured in the perioperative period and compared between patients with and without complications. Data are presented as mean ± SEM. Statistical analysis was performed using two-way analysis of variance (ANOVA) with multiple comparisons test and *post hoc* Sidak's correction. * denotes p<0.05.

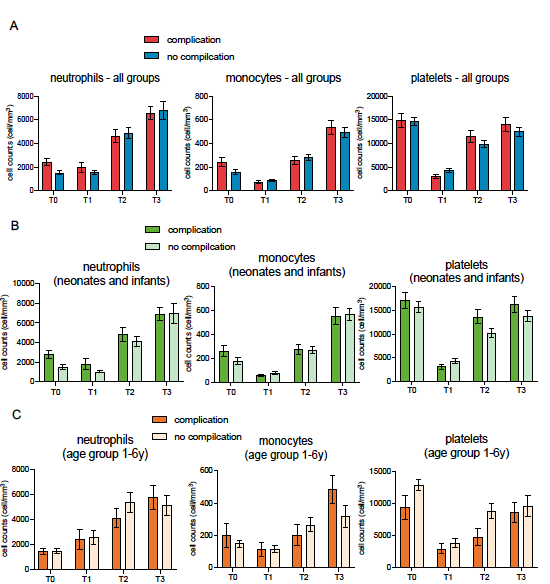
**Suppl Fig. 2 Comparison of neutrophil and platelet counts in patients with and without postoperative complications at different time points. (A)** Comparison of neutrophil, monocyte and platelet counts in the overall cohort, distinguishing patients with and without postoperative complications. Subgroup analysis in neonates and infants **(B)** and patients aged 1-6 years **(C)**, examining the association between postoperative complications and neutrophil and platelet counts. Data are presented as mean ± SEM. Statistical analysis was performed using two-way analysis of variance (ANOVA) with multiple comparisons test and *post hoc* Sidak's correction.

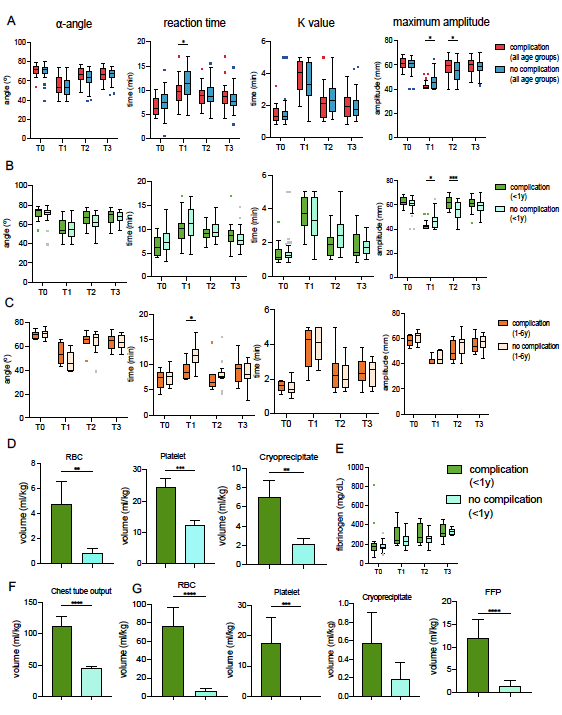

**Suppl Fig. 3 Comparison of TEG parameters in patients with and without postoperative complications at different time points. (A)** Comparison of TEG values, including ⍺-angle, reaction (R) time, k time, and maximum amplitude (MA) from T0 to T3 in the overall cohort, distinguishing patients with and without postoperative complications. Subgroup analysis in neonates and infants **(B)** and patients aged 1-6 years **(C)**, examining the association between postoperative complications and TEG values. Data are presented as median and interquartile range (IQR). (A-C) Statistical analysis was performed using two-way analysis of variance (ANOVA) with multiple comparisons test and *post hoc* Sidak's correction. * and *** denote p<0.05 and 0.001, respectively. **(D-G)** Comparison of intraoperative blood transfusion **(D)**, perioperative fibrinogen levels **(E)**, postoperative chest tube output **(F)**, and postoperative blood transfusion **(G)** between neonates/infants with complications and without complications. Data are presented as mean ± SEM. Statistical analysis was performed using Mann-Whitney test (D,F,G) and two-way analysis of variance (ANOVA) with multiple comparisons test and *post hoc* Sidak's correction (E). **, ***, and **** denote p<0.01, 0.001, and 0.0001, respectively.

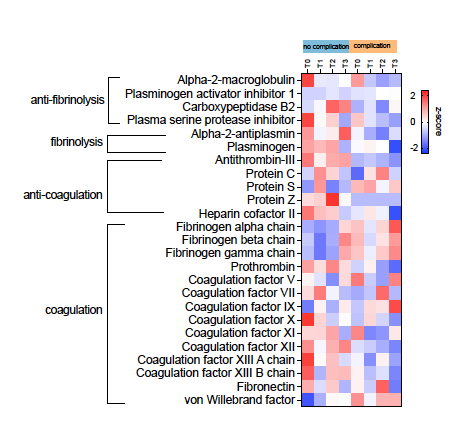
**Suppl. Fig. 4.** Heatmap illustrating the temporal changes in relative coagulation and fibrinolytic protein levels from T0 to T3 compared between patients with and without complications.

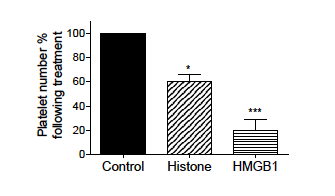

**Suppl. Fig. 5**. Platelet counts were compared before and after histone and HMGB1 stimulation. Data are presented as mean ± SEM. Statistical analysis was performed using one-way analysis of variance (ANOVA) with multiple comparisons test and *post hoc* Sidak’s correction. ** and *** denote p<0.01 and 0.001, respectively.

**Supplemental Table**

**Supplemental Table 1. MA cutoff for complications**

**ROC curve analysis for neonate/infant MA at T1 and postoperative complications**

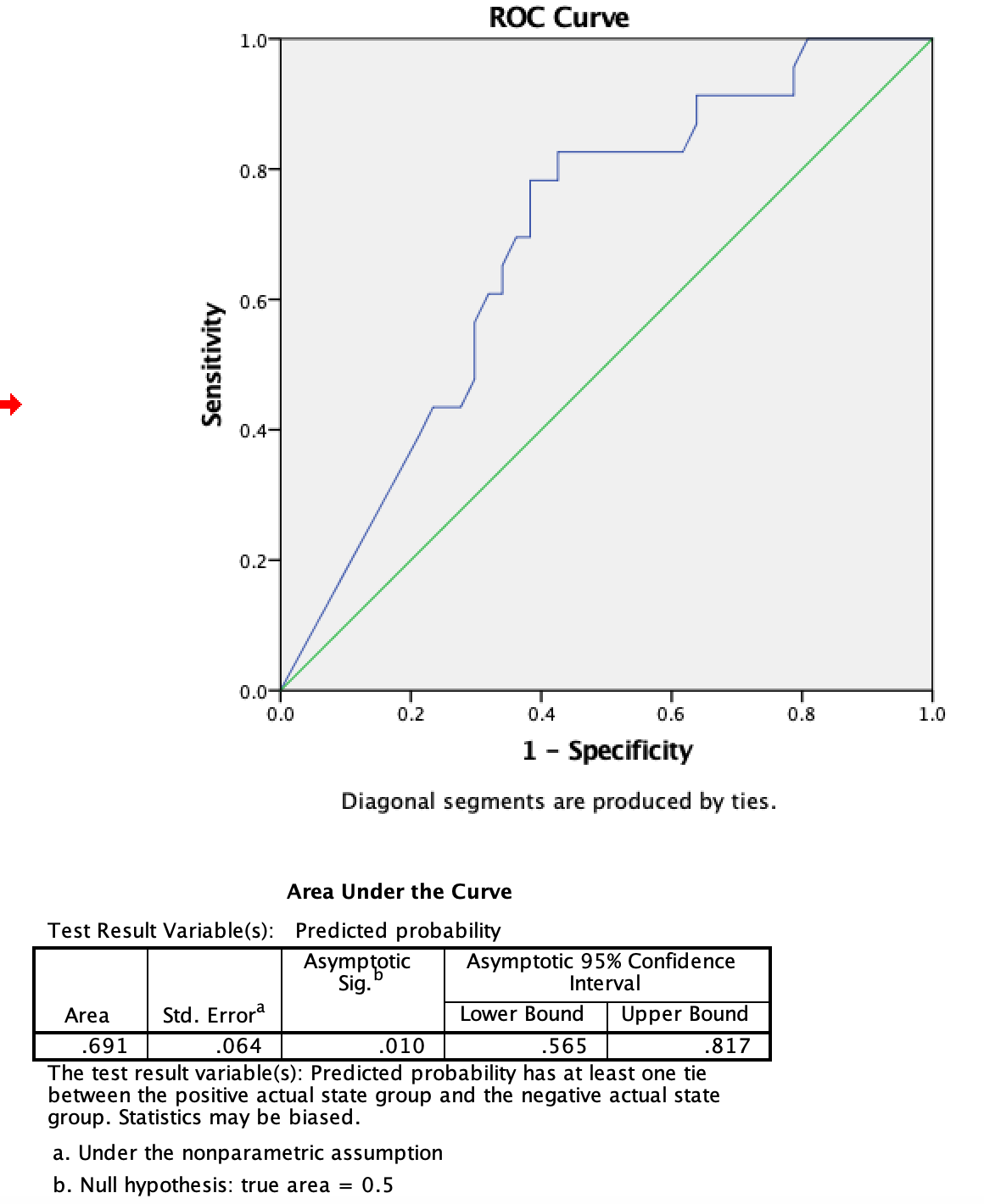

**Youden-J index is highest at MA of 44 mm (sensitivity = 82.6%, specificity = 57.4%)**

MA of <44 mm at rewarming is associated with postoperative complications
